# Deep proteome analysis of plasma reveals novel biomarkers of mild cognitive impairment and Alzheimer’s disease: A longitudinal study

**DOI:** 10.1101/2022.01.30.478370

**Authors:** Gurjeet Kaur, Anne Poljak, Perminder Sachdev

**Affiliations:** Centre for Healthy Brain Ageing, School of Psychiatry, University of New South Wales, Sydney, NSW, 2052, Australia; Mark Wainwright Analytical Centre, Bioanalytical Mass Spectrometry Facility, University of New South Wales, Sydney, NSW, 2052, Australia; Neuropsychiatric Institute, Euroa Centre, Prince of Wales Hospital, Sydney, NSW, 2052, Australia

**Keywords:** Plasma, proteomics, preclinical biomarkers, longitudinal study, ageing, mild cognitive impairment, Alzheimer’s disease

## Abstract

Ageing is the primary risk factor for AD; however, there is a poor understanding of the biological mechanisms by which the ageing process contributes to the development of AD in some individuals, while others progress to advanced age with relatively little AD neuropathology. To halt the progression of AD, the preclinical stage of neurodegeneration (before the onset of clinical symptoms) is anticipated to be the more effective time point for applying potentially disease-modifying interventions in AD. The main objective of this study was to understand the age and disease related proteomic changes are detectable in plasma, based on retrospective analysis of longitudinal data and cross-sectional analyses of clinically diagnosed cases. We conducted an in-depth plasma proteomics analysis using intensive depletion of high-abundant plasma proteins using the Agilent multiple affinity removal liquid chromatography (LC) column-Human 14 (Hu14) followed by sodium dodecyl sulfate-polyacrylamide gel electrophoresis (SDS PAGE) technique. In this study, we have begun to address the following questions; (1) differences in plasma proteomic profiles between normal ageing, vs ageing with progress to cognitive decline (MCI) or disease (dementia, probable AD), (2) cross-sectional analysis of baseline data, when all subjects are clinically identified as cognitively normal, provides insight into the preclinical changes which precede subsequent progression to AD and potentially provide early biomarkers, and (3) comparison of plasma at the point of progression to clinically diagnosed onset of cognitive decline or AD, can provide potential plasma biomarkers to facilitate clinical diagnosis. Furthermore, our findings also identified some proteins previously discovered in AD CSF and brain proteomics signatures that could provide clinically meaningful information. We identified differentially expressed proteins which were associated with several biological and molecular processes that may serve as therapeutic targets and fluid biomarkers for the disease.

## Introduction

Alzheimer disease (AD) accounts for up to 70% of all dementia cases and is the most common cause of dementia. Ageing is the primary risk factor for AD; however, there is a poor understanding of the biological mechanisms by which the ageing process contributes to AD development in some individuals, while others progress to advanced age with relatively little AD neuropathology. The pathogenesis of AD is now recognized to be multifactorial, with dysregulation of various cellular and molecular processes contributing to the disease process, including synaptic damage, mitochondrial dysfunction, and oxidative stress^1–6^. While advancing age is the single greatest risk factor for AD^7^, other factors such as *APOEε4* allele^8^, comorbidities such as vascular disease^9^ and lifestyle factors such as head injuries^10,11^ all contribute to the level of AD risk. Early AD manifests clinically as mild cognitive impairment (MCI)^12^, particularly in the case of amnestic MCI, although a clinical diagnosis of MCI often stays stable or even reverts to normal and does not always progress to dementia^13,14^. By the time AD manifests as dementia, the level of brain pathology may be impossible to revert since substantial neuronal cell death has occurred. Identifying biomarkers of transition from normal to MCI (if not earlier) might provide a window of opportunity for prevention trials that focus on ameliorating symptoms before neurodegeneration progresses to clinically identifiable symptoms. Several neuroimaging and CSF based biomarkers for diagnostic evaluation of dementia have recently been recommended by an international consensus group^15^. The major limitations with CSF and neuroimaging biomarkers are that they are not likely to be widely adopted for routine use or population screening due to their invasive nature, high cost, limited availability and requirement of high-level technical skill and training to implement^16^. By contrast, blood is a relatively easy fluid to collect, and venepuncture is a routine and commonly performed procedure for clinical and research purposes^16^.

Mass spectrometry-based methods represent the only unbiased approach for discovery focused proteome analysis. They are rapid, sensitive, can provide both qualitative and quantitative information, and for the study of proteins, can also provide information about post-translational modifications and protein interactions^17^. The main obstacle has been identifying methods of narrowing the extreme dynamic range of the plasma proteome while maintaining sufficient methodological simplicity to apply to moderately sized clinical studies^18^. Recent advances in plasma proteomics have identified promising approaches to achieve the depth of plasma proteome coverage using prefractionation methods^19–21^. In the current study, we used a two-step plasma fractionation approach; Agilent multiple affinity removal liquid chromatography (LC) column-Human 14 (Hu14) followed by sodium dodecyl sulfate-polyacrylamide gel electrophoresis (SDS PAGE) technique, based on our previously published method^20^. This workflow facilitated extended plasma proteome coverage unbiased, allowing identification of biologically meaningful longitudinal and crosssectional proteome changes in individuals progressing through stages of cognitive impairment over the decade in which greater risk commences. A set of potential protein biomarkers might facilitate the development of precise tests for detecting the disease at the early stages. Furthermore, these markers may help identify unexpected biological pathways and new potential therapeutic targets for future development.

## Materials and Methods

### Cohort, plasma and proteomics experimental design

Plasma samples were obtained from the Sydney Memory and Ageing Study (MAS) from participants aged 70-90 years^22^. The baseline sample was collected (Wave 1) between September 2005 and November 2007, at which time all participants were cognitively normal (n = 33). Participants were followed up for six years (Wave 4), with 11 participants remaining normal and the remainder progressing to MCI and AD (n=11 each). The diagnosis was by consensus and met the NIA-AA criteria for MCI and Alzheimer’s dementia, respectively (Table 1). Detailed inclusion and exclusion criteria for the MAS cohort was previously published^22^. We selected only individuals with aMCI (amnestic MCI) for this study, as this subtype is generally related to subsequent progression to Alzheimer’s dementia^23,24^. Blood was collected into EDTA containing tubes, centrifuged (2000*g*, 20min, 4°C), and the plasma transferred into clean 1.5 mL polypropylene tubes. To minimize freeze-thaw cycles, plasma aliquots were prepared (50, 250 and 500 μL) and stored at −80°C until required. The UNSW Human Research Ethics Committee approved a protocol for blood collection (MAS ethics number; HC14327).

**Table 1:** Details of the participant demographics which were included in our present study.

| Total participants | Normal ageing | MCI | AD | Kruskal-Wallis statistic | Kruskal-Wallis P value |
| --- | --- | --- | --- | --- | --- |
| Total participants in each wave | 11 | 11 | 11 | NA | NA |
| Wave 1 age in years mean±SD (CV%) | 76.89±3.39 (4.41%) | 78.46±5.5 (7.01%) | 80.62±4.72 (5.85%) | 4.32 | 0.11 |
| Wave 4 age in years mean±SD (CV%) | 82.95±3.35 (4.04%) | 84.37±5.59 (6.63%) | 86.51±4.81 (5.56%) | 4.38 | 0.11 |
| Education (years) at wave 1 | 10.84±4.05 (37.44%) | 10.77±3.92 (36.39%) | 10.48±2.53 (24.16%) | 0.04 | 0.97 |
| Length of follow up (years) | 5.77 | 5.95 | 5.75 | NA | NA |
| Clinical diagnosis at W1 | Normal | Normal | Normal | NA | NA |
| Clinical diagnosis at W4 | Normal | amdMCI | Dementia, probable AD | NA | NA |
| W1 APOE status | E3/3 | E3/3 | E3/3 | NA | NA |
| MMSE at W1 mean±SD (CV%) | 29.36±1.50 (5.11%) | 28.27±1.55 (5.50%) | 28.36±1.80 (6.36) | 5.61 | 0.06 |
| MMSE at W4 mean±SD (CV%) | 29.55±0.93 (3.16%) | 28.18±1.40 (4.97%) | 23.64±4.05 (17.16%) | 20.92 | 0.00 |
| Total WMH volume W1 mean±SD (CV%) | 16032±19357 (120.7%) | 9274±4112 (44.34%) | 12612±13117 (104%) | 0.28 | 0.87 |
| Total WMH volume W4 mean±SD (CV%) | 21407±14975 (69.96%) | 17145±7380 (43.05%) | 33005±20872 (63.24%) | 3.40 | 0.18 |
| BMI (median) mean±SD (CV%) | 26.91±4.98 (18.54%) | 28.73±6.05 (21.06%) | 26.82±2.89 (10.78%) | 1.01 | 0.60 |
| Cholesterol (mmol/L) | 5.08±0.98 | 4.41±0.75 | 4.79±0.72 | 3.41 | 0.18 |
| mean±SD (CV%) | (19.32%) | (17.11%) | (15.09%) |  |  |
| Triglyceride (mmol/L)<br>mean±SD (CV%) | 1.36±1.36<br>(99.76%) | 1.26±0.71<br>(56.88%) | 0.90±0.35<br>(39.75%) | 1.47 | 0.47 |
| HDL-Chol (mmol/L)<br>mean±SD (CV%) | 1.40±0.40<br>(28.99%) | 1.20±0.35<br>(29.81%) | 1.34±0.32<br>(24.05%) | 2.16 | 0.33 |
| LDL-Chol (mmol/L)<br>mean±SD (CV%) | 3.16±0.80<br>(25.46%) | 2.64±0.66<br>(25.04%) | 3.02±0.67<br>(22.31%) | 2.71 | 0.25 |
| Glucose (mmol/L)<br>mean±SD (CV%) | 6.58±2.76<br>(42.05%) | 6.31±1.13<br>(18%) | 5.80±0.58<br>(10.11%) | 1.43 | 0.48 |
| Urate (mmol/L)<br>mean±SD (CV%) | 0.32±0.07<br>(24.5%) | 0.35±0.04<br>(13%) | 0.35±0.08<br>(24.08%) | 1.40 | 0.49 |
| Vitamin A (umol/L)<br>mean±SD (CV%) | 2.80±0.48<br>(17.42%) | 3.51±0.21<br>(20.10%) | 3.15±1.27<br>(40.54%) | 5.81 | 0.05 |
| Vitamin E (umol/L)<br>mean±SD (CV%) | 44.13±33.77<br>(76.53%) | 31.89±6.40<br>(20.08%) | 31.74±9.14<br>(28.22%) | 2.18 | 0.33 |
| Carotene (umol/L)<br>mean±SD (CV%) | 1.03±0.76<br>(73.07%) | 0.63±0.39<br>(62.94%) | 0.79±0.52<br>(66.38%) | 1.53 | 0.46 |

SD= standard deviation, cv= coefficient of variations

Proteomics profiling was performed on 33 humans (66 total) plasma samples from wave 1 (baseline) and wave 4 (6 years follow up), in the following three groups: (1) individuals cognitively normal at wave 1 denoted as <u>CTRLW1</u>, who remained normal at wave 4 denoted as <u>CTRLW4</u>, (2) individuals cognitively normal at wave 1 denoted as <u>MCIW1</u> who progressed to MCI at wave 4 denoted as <u>MCIW4</u>, (3) individuals cognitively normal at wave 1 denoted as <u>ADW1</u> who progressed to dementia, probable AD at wave 4 denoted as <u>ADW4</u>.

### Depletion of high abundant proteins using Human 14 (HU14) immunoaffinitybased Columns

The protocol followed for plasma high abundance protein removal, and fractionation of the low abundance proteins, was adapted from a previously published approach^20^. The approach involved depletion of the top 14 high abundance proteins (albumin, IgG, antitrypsin, IgA, transferrin, haptoglobin, fibrinogen, α-2-macroglobulin, α-1-acid glycoprotein, IgM, apolipoprotein AI, apolipoprotein AII, complement C3, and transthyretin) using an Hu14 column (4.6 x 100 mm, Agilent California, United States), followed by SDS/PAGE fractionation of the low abundance protein fraction. We describe the approach here in brief; Plasma (50 μL) was diluted by adding 150 μL of Agilent buffer A (1:4 dilutions, as recommended by Agilent Technologies), then filtered using a 0.45μm spin filter (Spin-X centrifuge tube filter, 0.45 μm Cellulose Acetate, Merck, Germany) to remove particulates. Samples were then injected (100 μL) onto the Hu14 column. Chromatography and fraction collection were carried out on an Agilent 1290 UPLC system (Agilent, Santa Clara, CA), using Hu14 buffers A and B purchased from Agilent (Santa Clara, CA), following the manufacturer’s instructions (Agilent, Santa Clara, CA) for protein binding and elution. Only the low abundance protein fraction was further fractionated and analysed using LC-MS/MS.

### Fractionation of low abundance proteins using 1D-SDS PAGE, tryptic digest and LC-MS/MS

Equal amounts of total protein (50 μg) from the Hu14 depleted plasma were filtered using Amicon ultra 3kDa centrifugal filter units (MERCK, New Jersey, USA), dried in speed vac (ThermoFisher, Massachusetts, USA) and diluted to a final volume of 20 μL by adding 5 μL LDS sample buffer Invitrogen NuPAGE (ThermoFisher, Massachusetts, USA), 2 μL reducing agent Invitrogen NuPAGE (ThermoFisher, Massachusetts, USA), and 13 μL deionized water (MilliQ). Samples were then briefly heated (10 minutes, 70°C), followed by electrophoresis; 1D SDS/PAGE using Invitrogen NuPAGE 4-12% gradient Bis-Tris midi gels (ThermoFisher Scientific, Massachusetts, USA) and 1x Invitrogen MES running buffer according to the manufacturer’s instructions (ThermoFisher Scientific, Massachusetts, USA, USA) followed by colloidal coomassie G250 staining^25^ (Figure S4). After destaining, the separated protein lanes were cut evenly with a 24-lane blade (Gel Company Inc., CA), and the gel slices were collected into ten vials for destaining, in-gel trypsin digestion and label-free LCMSMS, following the approach taken in our previously published work^20^.

### Computational Analysis

Computational analysis of the raw files was performed for protein identification and quantification. The consistency of protein expression change was determined using two label-free quantification approaches, peak area integration and spectral counting. Protein identification, peak area integration and fold-change calculation were performed using ProteomeDiscoverer v2.4 software (Thermo Fisher Scientific, Waltham MA), in conjunction with three search engines (Mascot, Sequest, and Amanda). Protein identification followed by spectral counting and fold-change determination was carried out using a combination of Mascot search engine and Scaffold Q+ software v 4.11.0 (Proteome Software, Portland, OR). A minimum of ≥2 unique peptides per protein were required for protein identification and quantitation on all data analysis software. The UniProt *Homo sapiens* (human) database was combined with reversed decoy database to determine FDR by all search engines for MS and MS/MS spectral mapping. Mass tolerance for matching peaks to theoretical ion series was five ppm. False discovery rate (FDR) was set to <1% to ensure only high-confidence protein identifications. Enzyme specificity was set to trypsin, with a maximum of two missed cleavages. Searches included variable modifications of protein N-terminal acetylation, methionine oxidation, and fixed modification of carbamidomethylation of cysteines. All the parameters were kept similar in both search engines. To select only those proteins with robust expression change between groups, we used the following inclusion criteria: only those proteins quantified in >6 individuals, proteins identified with a minimum of two peptides per protein, the consistent direction of protein fold change across two bioinformatics platforms with orthogonal quantification approaches (peak area ratio with PD2.4 and spectral counting with Scaffold) with a fold change of at least 20% (≤0.08 and ≥ 1.2) in preferably both search engines but at least one. These orthogonal approaches have specific advantages and disadvantages^26,27^, so we reasoned that the most reliable changes should be consistent across platforms.

### Bioinformatics Analyses

We used RStudio version 1.2.5033 and R version 3.6.3 for most post data processing analyses, including heatmap and volcano plots. Venny 2.1 was used to plot Venn diagrams^28^. We performed gene ontology (GO) term enrichment analysis using differentially expressed proteins (DEPs) to compare biological processes and pathways affected in normal ageing, MCI and AD, using STRING (version 11.5). This kind of analysis uses GO terms to classify proteins into particular roles or functions (i.e., biological processes, cellular components, molecular function and KEGG & Reactome pathways). From this kind of sorting, we can identify numbers of proteins that subserve specific functions (i.e., “observed gene count” within the STRING output). Information about the level of enrichment of functional categories is also provided by comparison with a background set of proteins (we used the default whole human genome available within STRING for the analyses presented here), which allows an estimation of the enrichment score (strength) and level of statistical significance (FDR). Together the observed gene counts and enrichment strength values give an idea of which functional categories are represented by (a) the most significant number of proteins and (b) are most enriched relative to the background set. Both observations help identify functional categories that are associated with the disease. However, it should be noted that (1) most proteins are pleiotropic and may be listed within several functional groups, and (2) the GO term lists are a manually curated artificial construct and include some very broad terms which may capture many proteins (e.g., cellular process, biological regulation, binding, and others), but which are minimally informative from a specific function perspective. For this reason, observed gene count and enrichment strength values generally vary in an approximately reciprocal manner and therefore should be used together to identify biological/disease relevance functions. It is likely that categories of the greatest relevance will be those with a moderate score for both observed genes count and enrichment strength, rather than those that fall at the extremes of either value.

## Results

### Overview of proteomics study populations

The main objective of this study was to discover detailed plasma biomarker profiles reflecting normal ageing, mild cognitive impairment (MCI) and dementia, probable Alzheimer’s disease (AD). Participant demographics are shown in Table 1.

Since each plasma sample consisted of ten fractions, a total of 660 LC-MSMS runs were performed to maximize plasma proteome coverage of low abundant proteins. In total, we identified 1,578 proteins (false discovery rate <1%) with 32,469 total peptides using Proteome Discoverer 2.4 search engine. Data analysis was performed on 2 different search engines, i.e., Proteome Discoverer 2.4 and Scaffold Q+ software v 4.11.0. We performed analyses in six different combinations, including both longitudinal and cross-sectional analyses. Longitudinal analyses included: **1.** Ageing while maintaining normal cognition <u>(CTRLW4/CTRLW1)</u> **2.** MCI <u>(MCIW4/MCIW1)</u>; **3.** AD <u>(ADW4/ADW1)</u> and cross-sectional analyses included; **4.** MCI vs age-matched controls <u>(MCIW4/CTRLW4)</u>; **5.** Incipient AD vs age-matched controls <u>(ADW4/CTRLW4)</u>; and **6.** preclinical AD vs age-matched controls; <u>(ADW1/CTRLW1)</u>. The longitudinal analyses provide insight into changes that occur in normal ageing over 6 years and a progression from clinically normal to MCI or AD over 6 years of baseline to follow-up. These longitudinal analyses allow comparison of ageing while retaining clinically normal cognition and ageing with progression to cognitive disease and dementia, suggesting proteins and pathways which are disrupted in the development of disease/disorder. By contrast, the cross-sectional analyses compare incipient MCI or AD to cognitively normal age-matched controls, which may identify potential disease biomarkers. Figure 1A illustrates the samples and mass spectrometry method employed in this work. Venn diagrams showed an overlap of 1,467 proteins in the three longitudinal comparisons (Figure S1C) and 903 proteins in four cross-sectional comparisons (Figure S1D). Furthermore, we identified 450 differentially expressed proteins (DEPs) in the longitudinal comparisons and 553 DEPs in the crosssectional comparisons, summarized in Table S2a and S2b respectively. Scatter plots and density plots for all 6 comparisons are shown in Figures S2i and S2ii respectively. The detailed scatter plots were plotted using the DEPs from both the search engines i.e. PD2.4 and scaffold in all 6 comparisons Figure S2iii.

**Figure 1:**
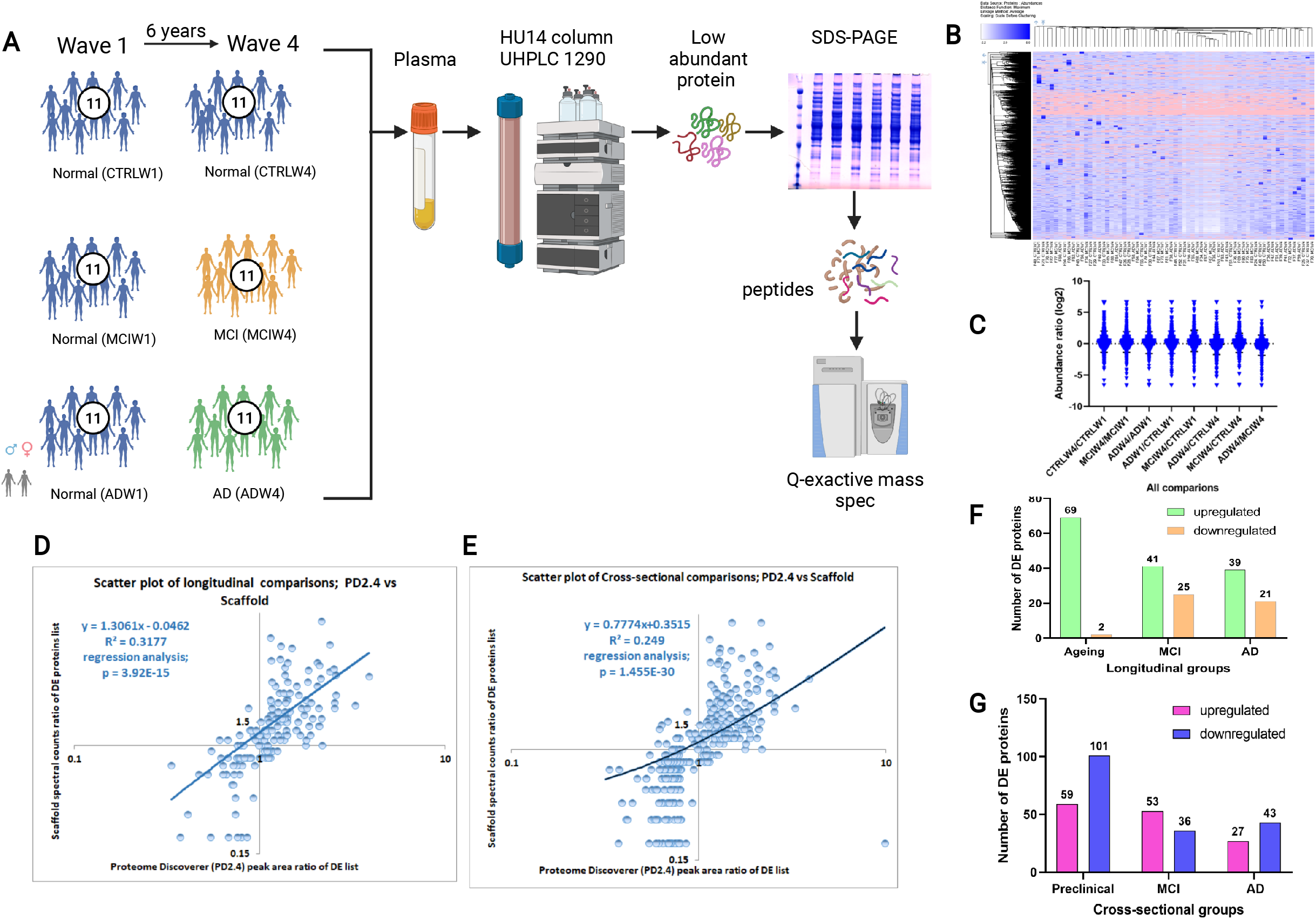
Proteome profiling and comparison of normal ageing, MCI, and AD in longitudinal and cross-sectional cohorts. **A.** Overview of the study population and schematic proteomic workflow. The plasma of two waves comprising ageing, MCI and AD subjects was analysed. The total number of subjects per group is depicted. Blue figurines represent cognitively normal individuals (regardless of wave), while orange and green figurines depict progression to MCI and AD respectively at W4, from normal cognition at W1. **B.** Hierarchical cluster analysis and heat map for 1,578 total proteins identified in 66 individual samples (output from ProteomeDiscoverer 2.4 (PD2.4) software). **C.** Scatter dot plot analysis using abundance ratios of all 6 comparisons used in this study, and providing a global view of level and direction of fold-change across comparison groups. Small horizontal lines, around the center of each cluster, show the mean and the error bars ± SD. **D and E.** Scatter plots and regression analysis comparing differentially expressed proteins (DEPs) identified in the two orthogonal quantitative methods; peak area ratio (PD2.4) and spectral counting (Mascot & Scaffold). The final list of DEPs identified by both quantitative approaches, and used in all longitudinal (D) and cross sectional (E) comparisons were used in the scatterplots. **F**. Global analyses of proteomic changes in longitudinal groups. Bar graph showing the total number of proteins upregulated (green bars) and downregulated (yellow bars) in normal ageing (CTRLW4/CTRLW1), MCI (MCIW4/MCIW1), and AD (ADW4/ADW1). The number of DEPs in each group are indicated at the top of each bar. **G.** Global analyses of proteomic changes in cross-sectional analysis groups. Bar graph showing the total number of proteins upregulated (pink) and downregulated (blue) in Preclinical AD (ADW1/CTRLW1), MCI (MCIW4/CTRLW4), and AD (ADW4/CTRLW4). The number of DEPs in each group are indicated at the top of each bar.

The plasma proteomes of 33 individuals (11 individuals in each category; normal control, MCI, and AD) are compared by hierarchical clustering analysis (HCA) (Figure 1B), abundance ratios (Figure 1C) and box and whisker plot (Figure S1A), showing very similar distribution patterns overall. This is expected since most identified protein’s expression is unaltered between samples, even in disease. Similarity matrix analyses (Figure S1B) show a close association of protein expression data between the following group ratios: CTRLW4/CTRLW1, MCIW4/MCIW1, ADW4/ADW1 and ADW4/CTRLW4, and MCIW4/CTRLW4. The two orthogonal methods of identifying differentially expressed proteins were compared using scatter plots and regression analyses (Figure 1D and 1E), showing significant regression between the two quantitative approaches (scaffold spectral counting and PD2.4 peak area integration). Bar graphs of the total number of proteins up and downregulated in longitudinal and cross-sectional comparisons are shown in Figure 1F and 1G. In all longitudinal comparison groups, more proteins are upregulated than downregulated; this difference is particularly pronounced in the normal ageing group, with 69 upregulated and only 2 downregulated proteins (Figure 1F). The numbers of proteins up and downregulated with age (over the 6 years of the longitudinal analysis) were similar in MCI and AD (Figure 1F). In the cross-sectional comparison groups, the number of up and downregulated proteins varies across groups (Figure 1G), with MCI and AD having similar total numbers of DEPs. Interestingly, the preclinical AD group (Figure 1G) had the greatest number of total DEPs, and also the more significant number of upregulated (59) and downregulated (101) proteins than either the incident clinical MCI or AD group.

### DEPs identified in longitudinal analyses of ageing regardless of diagnosis

Comparing proteomic expression differences across the longitudinal cohorts provides insight into age-related changes, which are common across all three clinical groups, and appear to be largely independent of diagnosis. We observed that 71 proteins were dysregulated with ageing, the majority of which were upregulated (Figure 1F, Figure S5 and Table S3). These 71 age-related DEPs were manually grouped into 12 protein functional categories based on gene ontology (GO) using the PD2.4 analysis outcomes (Figure S3A). The three functional groups with the highest number of age-related DEPs were cell signalling (35%), cytoskeleton and microtubules function (17%), and metabolism (15%) (Figure S3A). A variety of other categories represented ≤8% of total DEPs each (Figure S3A). Of 71 DEPs in normal ageing, only two proteins were decreased, these being methanethiol oxidase (SELENBP1) and neuronal adhesion molecule 1 (NCAM1) Figure S5. By contrast, proteins associated with inflammation (S100A9, S100A4, YWHA/14-3-3 family proteins), metabolic proteins (LDHA, LDHB, PKM, NME2), proteasome subunits (PSMA4, PSME2, PSMB8, PSMA6, PSMA5), and DNA binding and repair (ENO1, PARK7, CALR) were increased with ageing in all three clinical groups in the longitudinal analysis. However, they were not specific to disease (Figure S5 and Table S3). The complete list of proteins that are differentially expressed in ageing is shown in Table S3 and heatmap (Figure S5A), while a volcano plot shows the top 20 age-related DEPs with the greatest fold change (FC) (Figure S5B).

Several age-related DEPs with the highest fold change include the following: tropomyosin alpha-4 (TPM4, FC= 3.4, p=0.00), chloride intracellular channel protein 1 (CLIC1, FC=3.391, p= 0.00), rho GDP-dissociation inhibitor 2 (ARHGDIB, FC=3.126, p=0.00), 14-3-3 protein zeta/delta (YWHAZ, FC=3.09, p= 0.00), 6-phosphogluconolactonase (PGLS), nucleoside diphosphate kinase B (NME2, FC=8.5, p=0.04), and NCAM1 (FC= 0.2, p=0.00) (see Table S3 for the full list). GO enrichment analysis of these 71 DEPs was performed to understand the molecular pathways affected in normal ageing (Table S3) using the STRING bioinformatics tool, and enrichment in multiple GO-based categories was observed, including; 157 biological processes, 36 cellular components, 16 molecular functions, 48 KEGG and Reactome pathways (Table S4).

### DEPs that change longitudinally with progression to MCI and AD from normal cognition

A total of 60 DEPs were identified uniquely in the longitudinal AD group (progression from cognitively normal at W1 to AD at W4, 6 years later); 39 upregulated and 21 downregulated (Figure 1F and Table 5a). In longitudinal MCI, a total of 66 proteins were differentially expressed, with 41 upregulated and 25 downregulated (Figure 1F, Table 6a). Heatmaps based on differential protein abundance values from both search engines depicted overall reproducibility as well as individual protein expression profiles in AD and MCI in Figures 2A and 2C, respectively. Volcano plots highlight the top 20 DEPs with the greatest magnitude of longitudinal fold-change in AD and MCI (Figures 2B and 2D, respectively). Several DEPs unique to AD progression in W4, and that were significantly (p ≤0.05) upregulated include: tropomyosin alpha-1 chain (TPM1, fold change (FC=19.4; p=0.00), calpain small subunit 1 (CAPNS1, FC=2.6; p=0.00), caveolae-associated protein 2 (SDPR, FC=18; p=0.05), endoplasmin (HSP90B1, FC=1.5, p=0.01). Additionally, proteins that were significantly downregulated included: alpha-mannosidase 2x (MAN2A2, FC=0.56; p=0.02), olfactomedin-like protein 3 (OLFML3, FC=0.42; p=0.03), keratin, type II cuticular Hb6 (KRT86, FC=0.51; p=0.00), and serotransferrin (TF, FC=0.51; p=0.00). The complete list of DEPs unique to longitudinal AD group, is shown in Table S5a. Functional categories with the greatest proportional change relative to either ageing or MCI, and with DEPs unique to longitudinal progression to AD, were associated with metabolism (26%,), membrane trafficking (10%) and neuron & synapse (4%), all higher in AD than either control or MCI. In comparison, cell signalling (12%), cell adhesion (3%) and protein turnover (2%) are all lower in AD than either control or MCI (Figure S3). The presence of proteins in plasma belonging to this AD-progression specific groups implies functional disruptions which may have contributed to the progression of AD (Figure S3C). Two functional categories which were proportionately increased in both MCI and AD, relative to normal ageing were growth factors and extracellular functions (Figure S3). Their difference to normal ageing and common MCI and AD, suggests a possible association with cognitive impairment.

**Figure 2:**
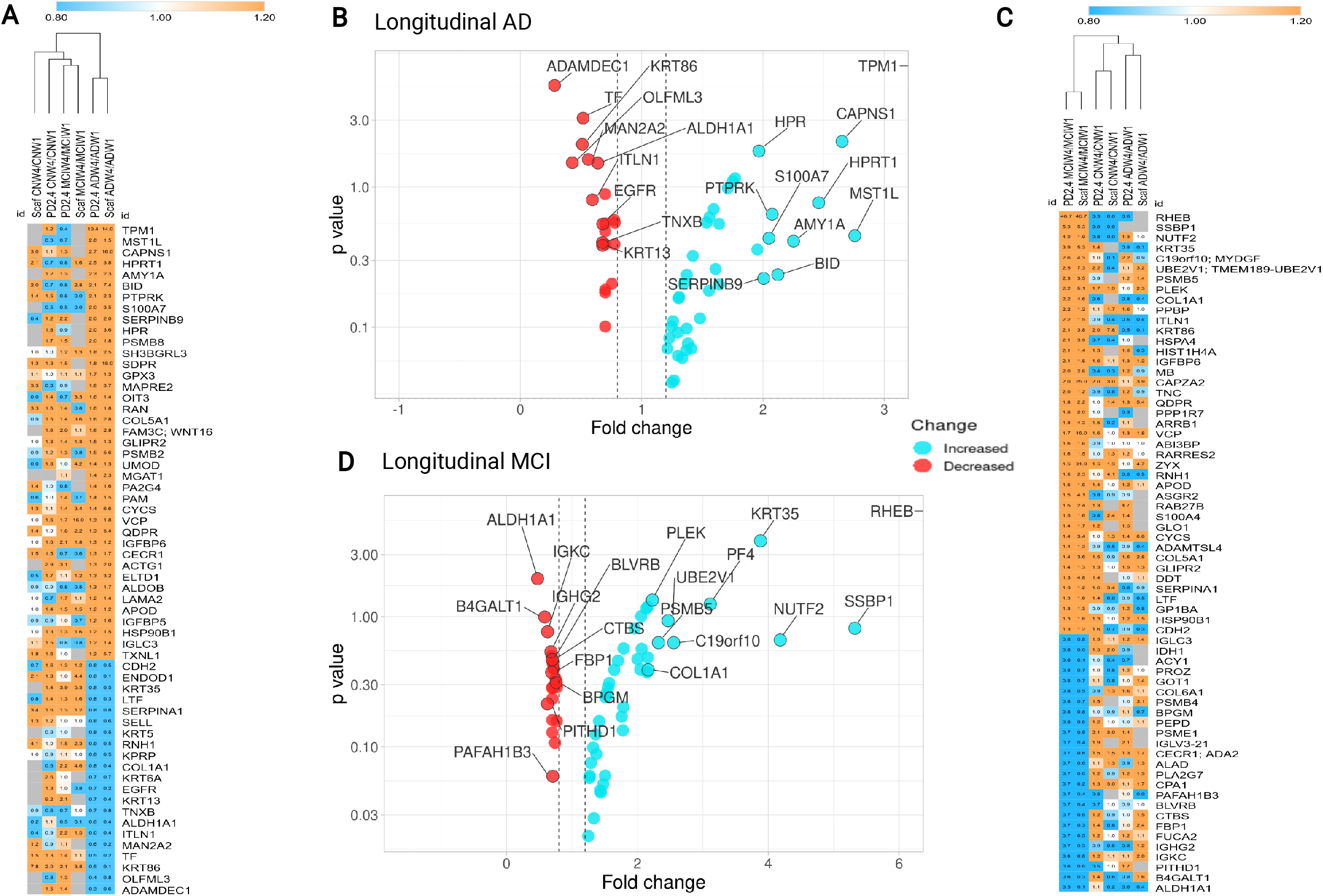
**A and C.** Heat map analysis of unique DEPs in longitudinal AD (ADW4/ADW1) and MCI (MCIW4/MCIW1) respectively. **B and D:** Volcano plots highlight the 20 DEPs with highest fold change in both the AD and MCI longitudinal analyses. The full list of DEPs, including p values, are shown in Tables S5a and Table S6a respectively.

In AD (Table S5a DEP list), 39 upregulated proteins were associated with 36 biological processes, 19 cellular components, 12 molecular functions, and 7 KEGG & Reactome pathways (Table S7). Approximately, half of the proteins were linked to binding activity (protein binding, signaling receptor binding, and calcium ion binding), stress response, small molecule metabolic process, extracellular regions, and cytoplasm.

The plasma proteome profile of longitudinal progression to MCI contained several unique DEPs not shared by AD and normal ageing W4 vs W1 groups (Figure 2C and 2D). In particular, the protein turnover group was proportionately higher in MCI (15%) than either the normal ageing group or AD, while cytoskeletal & microtubule structure was lower in MCI (4%) than either of the other groups (Figure S3B). MCI-specific DEPs with particularly high fold change with ageing, are shown in the Figure 2C and 2D heatmap and volcano plot and include upregulation of GTP-binding protein (RHEB, FC=46.691; p=0.00), pleckstrin (FC= 5.1; p=0.00), F-actin-capping protein subunit alpha-2 (CAPZA2, FC=25; p=0.00), insulin-like growth factor-binding protein 6 (IGFBP6, FC=1.8, p=0.01). Significantly downregulated proteins in MCI W4/W1 included flavin reductase NADPH (BLVRB, FC=0.4; p=0.02). The full list of DEPs in longitudinal MCI is shown in Tables 2 and S6a.

**Table 2:**
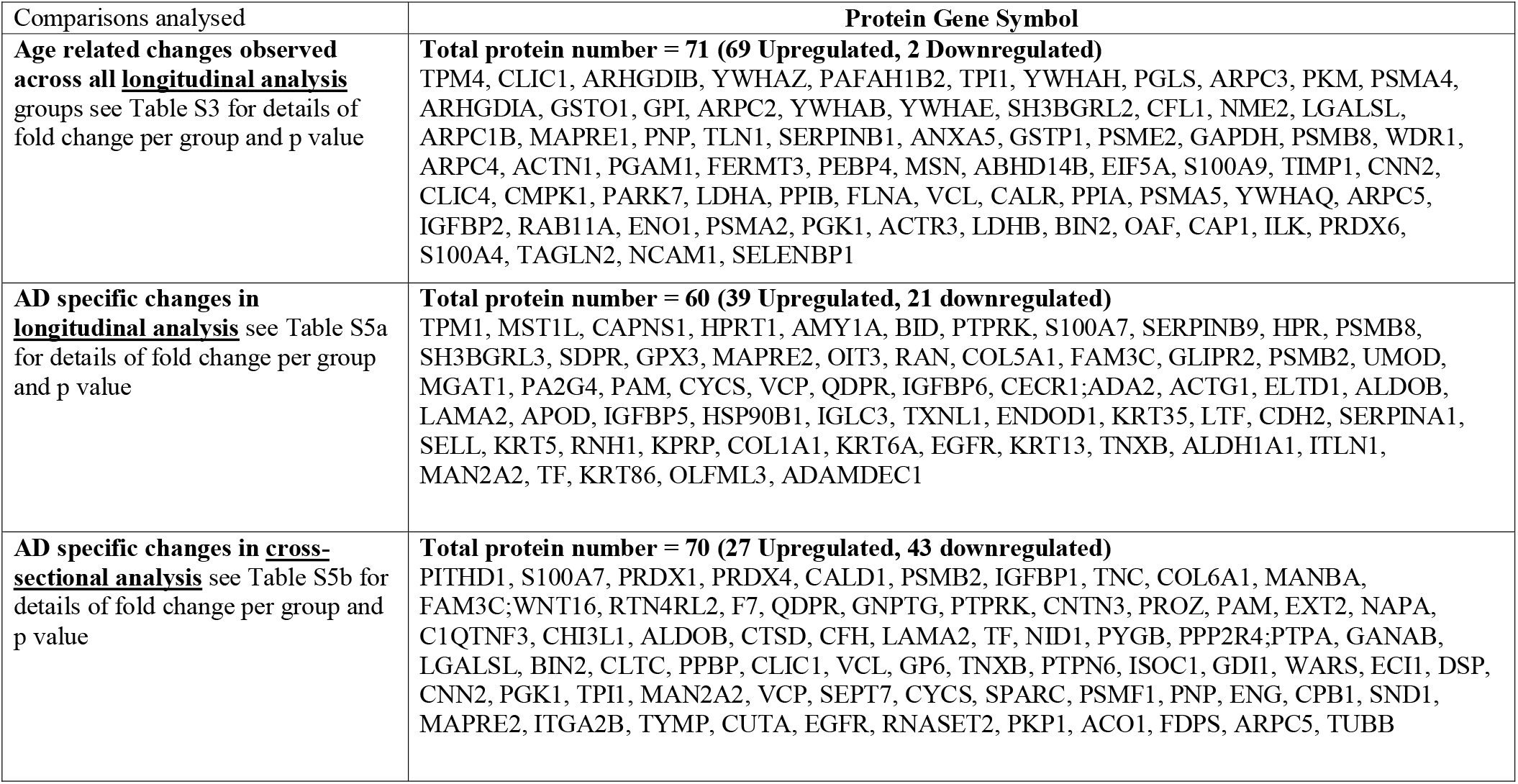
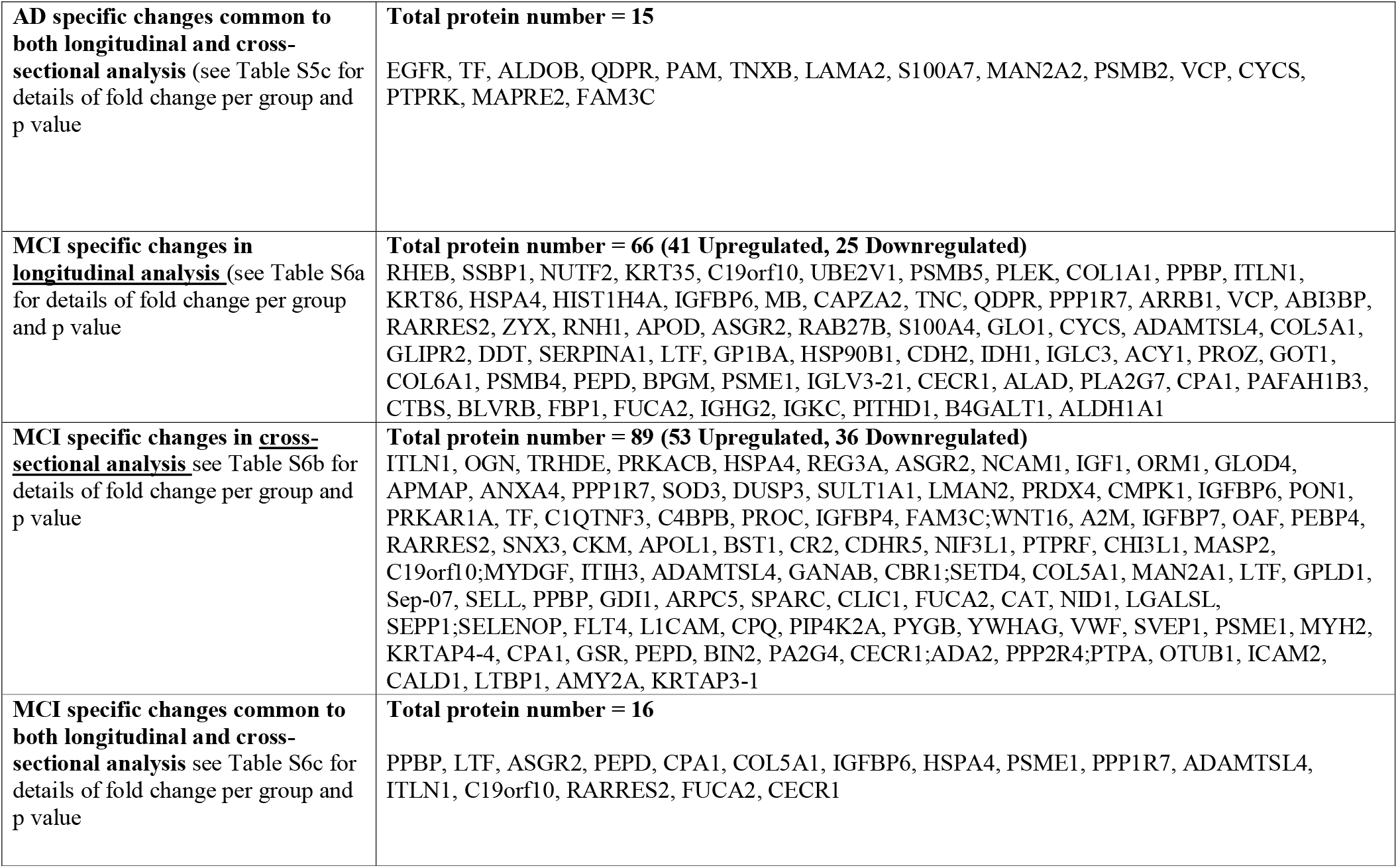
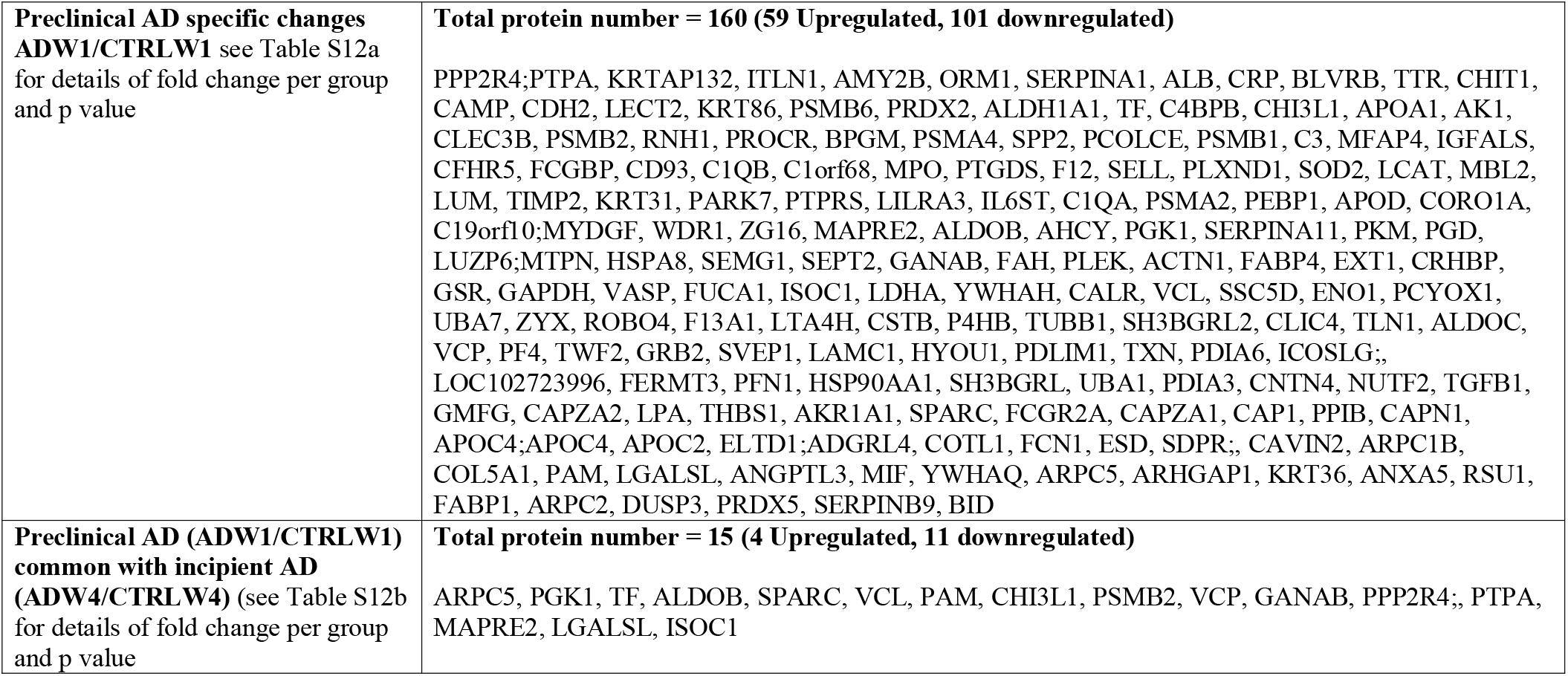
Summary table containing the final list of differentially expressed proteins (DEPs) in all the longitudinal and cross sectional comparisons analysed. This list contains DEPs quantified in >6 individuals per group, proteins identified with a minimum of two peptides per protein, consistent direction of protein fold change across two bioinformatics platforms with orthogonal quantification approaches (peak area ratio with PD2.4 and spectral counting with Scaffold) with a fold change of at least 20% (≤0.08 and ≥ 1.2) in preferably both search engines.

In MCI, upregulated proteins from Table S6a were based on GO term enrichment analysis were significantly associated with 99 biological processes, 22 cellular components, 7 molecular functions, and 2 KEGG & Reactome pathways (Table S8). Most DEPs fell into GO categories of; cellular process, cellular protein metabolism, regulation of protein phosphorylation, unfolded proteins, phosphate metabolic process, endomembrane system, signalling receptor binding and hemostasis. On the other hand, downregulated proteins from the MCI longitudinal analysis from Table S6a were significantly enriched in 33 biological processes, 10 molecular functions, and 11 KEGG & Reactome pathways (Table S8).

### Common plasma proteome changes in longitudinal AD and MCI groups

Only about 20% of total DEPs (19 DEPs) in AD and MCI longitudinal analysis groups were identified in both groups (Figure 3A). Of these 19 DEPs, 9 have the same direction of fold change; 8 are upregulated, and 1 are downregulated Figure 3B. The other 10 DEPs changed in the opposite direction in MCI and AD, showing that at the molecular level, substantial differences are apparent between MCI and AD, in that not only are a majority of DEPs different between the two groups (Figure 3A and 3B) but that even a good proportion of the proteins identified in common in the two groups have different directions of change.

**Figure 3:**
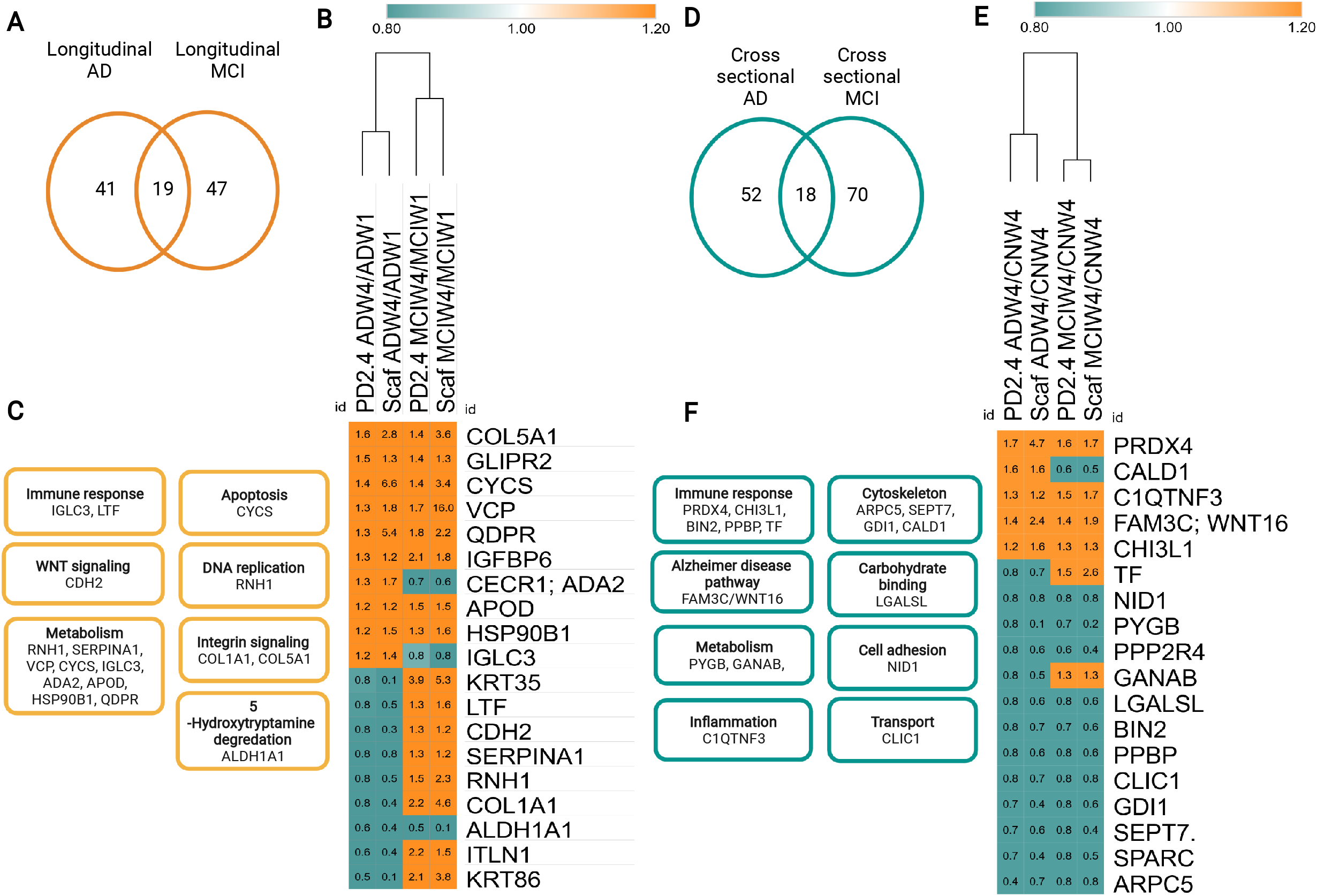
**A.** Venn diagram showing 19 DEPs that were present in both the longitudinal AD and MCI plasma proteome profiles. **B.** Heat map analysis of 19 DEPs common in longitudinal AD and MCI showing the pattern of common DEPs in both conditions. **C.** The 19 DEPs which were commonly dysregulated in MCI and AD were categorized into GO enrichment terms including metabolism, immune response, apoptosis, WNT signaling, DNA replication, Integrin signaling, 5-Hydroxytryptamine degradation, were associated with the list of MCI and AD common DEPs. **D.** Venn diagram showing 18 DEPs that were present in both cross-sectional AD and MCI plasma proteome profiles (Table S10b). **E.** Heat map analysis of 18 DEPs common to cross-sectional analyses of AD and MCI showing the expression pattern of common DEPs in both conditions. **F.** The 18 DEPs common to both AD and MCI cross-sectional analyses were categorized into 8 GO enrichment terms, including; immune response, cytoskeleton, Alzheimer’s disease pathways, protein folding, metabolism, cell adhesion, inflammation, ion transport and carbohydrate binding.

GO term enrichment analysis of the 19 DEPs shared by AD and MCI groups identified various functional groups, including metabolism, immune response, apoptosis, WNT signalling, and inflammation (Figure 3C). The two functional groups which have the greatest number of DEPs shared by both MCI and AD are metabolism, and immune response, suggesting that dysregulation of these two functions are shared between MCI and AD, while the majority of other DEPs are unique to each group (Figure 3A, 3B and 3C).

### Cross sectional proteome changes in AD and MCI – potential clinical biomarkers

Cross-sectional analyses compare the plasma proteome profiles of incipient AD and MCI relative to their age-matched cognitively healthy controls. In the cross-sectional analysis of incipient AD group (ADW4/CTRLW4), 70 DEPs were identified, including 27 upregulated and 43 downregulated DEPs (Table S5b). In MCI, 89 proteins were differentially expressed relative to age-matched normal controls (MCIW4/CTRLW4), with 53 upregulated and 36 downregulated DEPs (Table S6b), indicating that the number of dysregulated proteins identified in both disease conditions are similar. Heatmap analysis of the differentially abundant proteins in AD and MCI (Figure 4A and 4C respectively) show that there is some overlap of AD and MCI DEPs (Figure 3D). Volcano plots show the 20 DEPs in AD and MCI with the highest and lowest fold change in Figures 4B and 4D. Cross-sectional analyses of plasma proteome profiles of AD and MCI subjects relative to age-matched normal controls also identified a variety of potential disease-specific markers. DEPs identified in AD (ADW4/CTRLW4) that were not found in MCI (MCIW4/CTRLW4) (Figure 4A and 4B), including functions such as; antioxidants (PRDX4), proteasome (PSMB2, PSMF1), metabolism (MANBA, PYGB), cytoskeleton (TUBB, ARPC5). GO term enrichment analysis identified a diversity of significantly enriched categories in the DEPs upregulated in AD (9 biological processes, 14 cellular components, 7 molecular functions, 3 KEGG & Reactome pathways) and DEPs downregulated in AD (48 biological processes, 48 cellular components, 14 molecular functions, 2 KEGG & Reactome pathways), Table S10. Approximately half of the proteins were associated with binding activity (protein binding, signalling receptor binding and calcium ion binding), response to stress, small molecule metabolic process, extracellular regions, and cytoplasm.

**Figure 4:**
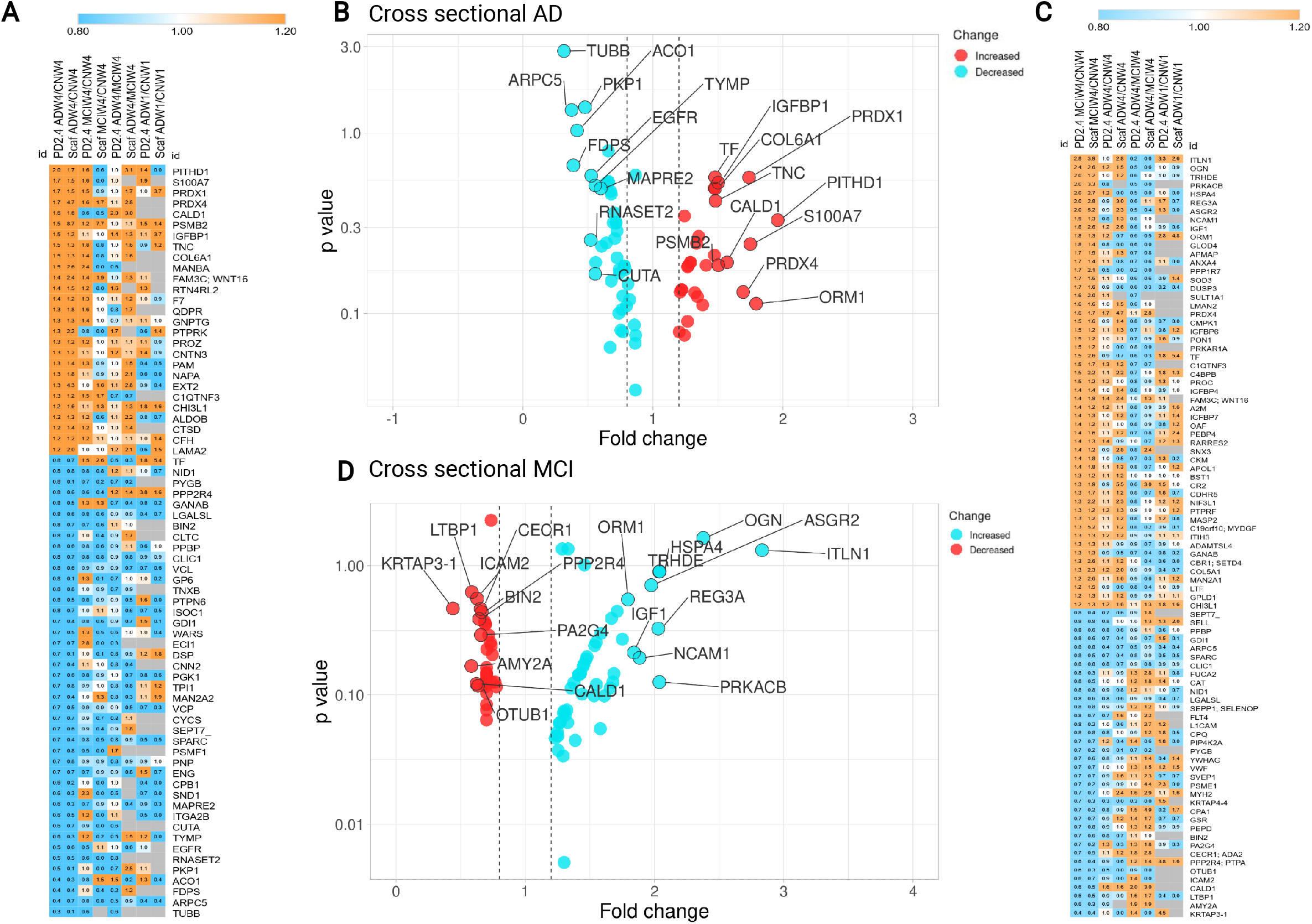
**A and C** Heat map analysis of DEPs in cross sectional comparisons of AD (ADW4/CTRLW4) and MCI (MCIW4/CTRLW4) respectively. **B and D:** Volcano plots highlight the 20 DEPs with highest fold change in cross sectional AD and MCI comparisons. The full list of DEPs, along with p values, are shown in Table S5b (AD) and Table S6b (MCI) respectively.

When compared to age-matched cognitively normal controls, the plasma proteome profile of MCI (MCIW4/CTRLW4) demonstrated a plethora of DEPs that were not observed in cross-sectional AD (ADW4/CTRLW4) group. These DEPs include functions such as growth factors (IGF1, MYDGF, OGN), metabolism (CBR1, GSR), signalling (YWHAG), immunity (ITLN1), and vascular function (VWF) (Figure 4C and 4D, Table S6b). GO term enrichment analysis identified 76 biological processes, 27 cellular components, 11 molecular functions and 6 KEGG & Reactome pathway categories significantly enriched (Benjamini-Hochberg FDR <0.05) using DEPs upregulated in MCI (Table S11). The main functional enrichments identified using the DEPs upregulated in MCI included: metabolic process, vesicle-mediated transport, immune system process, homeostasis and the complement cascade (Table S11). Enrichment analysis of DEPs downregulated in MCI identified 9 biological processes, 20 cellular components, 6 molecular functions, and 2 KEGG & Reactome pathway categories (Table S11).

Relatively few of the same DEPs were identified in both the cross-sectional and longitudinal analyses of AD and MCI, being 15 (Figure S6A and S6B) and 16 (Figure S6C and S6D) DEPs, respectively.

### Common proteome changes in cross sectional analysis of AD and MCI

There were 18 DEPs common to both AD and MCI in the cross-sectional analyses depicted in Figure 3D, E, and F. These MCI and AD shared DEPs were associated with functions such as immune system (PRDX4, CHI3L1, BIN2, PPBP, TF), cytoskeleton (ARPC5, SEPT7, GDI1, CALD1) and metabolism (PYGB, GANAB). Only 4 of these upregulated DEPs had a similar direction of fold-change in both AD and MCI (PRDX4, CHI3L1, FAM3C, C1QTNF3), while opposite directions of fold change were observed for TF, GANAB, and CALD1, Figure 3E. The majority of DEPs common to both MCI and AD in the crosssectional analyses were downregulated (11/18 proteins), Figure 3E.

Relatively few of the same DEPs were identified in both the cross-sectional and longitudinal analyses of AD and MCI, being 15 (Figure S6A, S6B and Table S5c) and 16 (Figure S6C, S6D and Table S6c) DEPs respectively.

### Plasma proteome changes in preclinical AD– potential early biomarkers

A retrospective analysis of baseline data allows us to identify potential early biomarkers of AD (i.e., ADW1/CTRLW1 ratios). We identified a total of 160 dysregulated proteins (Figure 5 and Table S12a), including 59 upregulated and 101 downregulated proteins in preclinical AD (ADW1/CTRLW1). The volcano plot and heatmap of all AD preclinical DEPs are depicted in Figure 5A and 5B. The volcano plot shows the top 20 most upregulated and downregulated DEPs (Figure 5A) and include functions such as metabolism (AMY2B, BLVRB), regulation (PPP2R4, SERPINA1), cytoskeleton (KRTAP13-2), immunity (ITLN1, RNH1, CRP, CHIT1), and transport (ALB, TTR), antioxidant (PRDX5), apoptosis (BID), signalling/regulatory (DUSP3, RSU1, ARHGAP1, YWHAQ, PAM), cytoskeleton (ARPC2), metabolism (FABP1), and anticoagulant (ANXA5) Figure 5A, Table S12 and S13.

**Figure 5:**
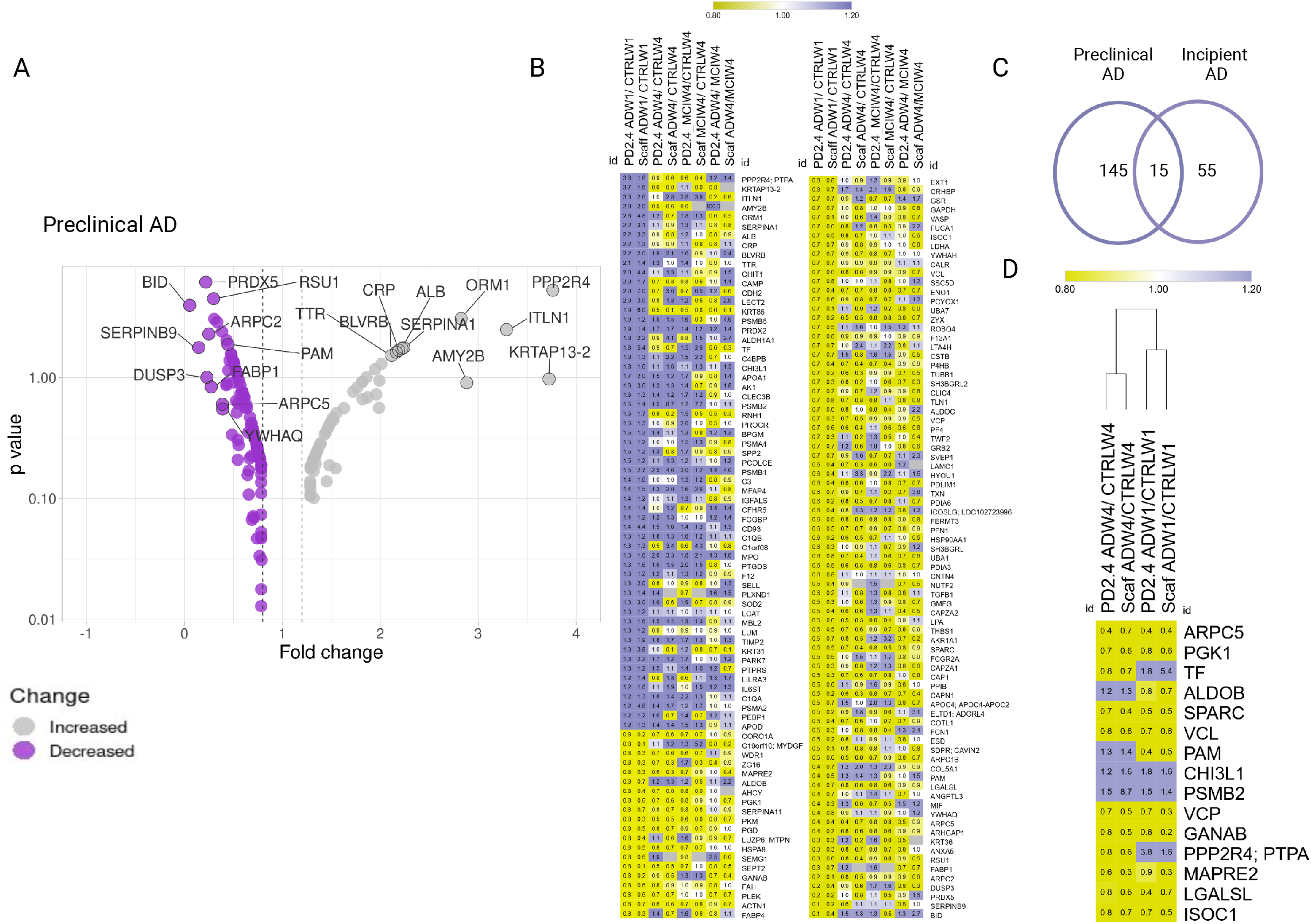
**A.** We identified a total 160 dysregulated proteins in preclinical AD (Figure 1G, and Table S12) including 59 upregulated and 101 downregulated proteins. The volcano plot highlights the 20 DEPs with highest fold change in preclinical AD (ADW1/CTRLW1) comparisons. Table S12 contains the full list and detailed information on the160 DEPs in preclinical AD. **B.** Heatmap of the 160 dysregulated proteins in preclinical AD. The 160 DEPs are presented in two panels for better visibility of fold change and protein names. **C.** Venn diagram showing the number of DEPs (15) which were common in preclinical AD (ADW1/CTRLW1). D. Common 15 DEPs are further presented in heatmap format.

A total of 15 DEPs were common to both incipient AD (ADW4/CTRLW4) and preclinical AD (ADW1/CTRLW1) (Figure 5C and 5D and Table 2, Table S12b). Of these, 9 DEPs were decreased in both clinical and preclinical AD, and included functions such as cytoskeleton/microtubule assembly (ARPC5, MAPRE2), signalling/regulation (PGK1), extracellular matrix (SPARC, VCL), apoptosis (VCP, ISOC1), protein folding (GANAB), and unknown (LGALSL). Two DEPs were increased in both preclinical and incipient AD: chitinase-3-like protein 1 (CHI3L1) and proteasome subunit beta type-2 (PSMB2). In addition, two DEPs were increased in preclinical AD but decreased in incipient AD: serotransferrin (TF) and serine/threonine-protein phosphatase 2A (PPP2R4), and two proteins were decreased in preclinical AD and increased in incipient AD; fructose-bisphosphate aldolase B (ALDOB) and peptidyl glycine alpha-amidating monooxygenase (PAM). These 15 DEPs may be potential preclinical plasma biomarkers of early AD.

In preclinical AD, 59 upregulated proteins were associated with 93 biological processes, 18 cellular components, 2 molecular functions, and 22 KEGG & Reactome pathways with significant GO enrichments Table S13. These GO-term enrichments were complement activation, post-translational protein modification, inflammatory response, neutrophil degranulation, metabolism, proteasome, and immune system. The 101 downregulated proteins involved 143 biological processes, 42 cellular components, 26 molecular functions, and 35 KEGG. Most DEPs were found to be involved in one of the following categories: immune system, actin cytoskeleton and polymerization, response to unfolded proteins, protein binding, glycolysis/gluconeogenesis, signalling by rho GTPases and haemostasis. Table S13 contains all the enriched GO with extensive information collected from STRING software.

## Discussion

This study has shown that a rich abundance of age and disease-related proteomic changes are detectable in plasma, based on retrospective analysis of longitudinal data and cross-sectional analyses of clinically diagnosed cases. In combination with a method which provides the depth of plasma proteome coverage^20^, the study design has addressed the following questions: (1) differences in plasma proteomic profiles between normal ageing and ageing with progression to cognitive decline (MCI) or AD; (2) cross-sectional analysis of baseline data, when all subjects were clinically identified as cognitively normal, provides insight into the preclinical changes which precede subsequent progression to AD, and potentially provide early biomarkers; and (3) comparison of plasma at the point of progression to the clinically diagnosed onset of cognitive decline or AD, can provide potential plasma biomarkers to facilitate clinical diagnosis. We perceive two major obstacles in identifying plasma protein biomarkers for the common age-related neurodegenerative diseases: (1) the restricted current level of information regarding the plasma proteome longitudinal changes in normal vs diseased individuals, and (2) the even more limited knowledge of preclinical AD plasma proteome. We have begun to address both of these deficiencies in this work.

### Plasma proteome changes in ageing using a longitudinal analysis

Ageing is the primary factor associated with organ function decline, including age-related cognitive decline, and is a major risk factor for neurodegenerative diseases such as AD^29^. Consequently, it is common to use age-matched controls to study such disorders and diseases. However, the extent of change in the ageing plasma proteome, irrespective of disease, is less clear. Here we identified 71 proteins that were dysregulated during normal ageing, with the majority being increased. These proteins were identified in all three of our longitudinal groups (normal controls, MCI and AD, Table 2) and with a similar fold-change direction (Table S3). The hippo signalling pathway was particularly enriched with ageing (Table S4). This signalling pathway included DEPs of the 14-3-3 protein family (YWHAZ, YWHAH, YWHAE, YWHAB, YWHAQ, YWHAB) and actin gamma 1 (ACTG1), which were all upregulated (Table S3). Recent evidence suggests that the hippo signalling pathway is involved in neuroinflammation, neuronal cell differentiation, and neuronal death^30^. The 14-3-3 protein family is highly expressed in the brain and influences many aspects of brain function through interactions with a diverse set of binding partners, including neural signalling, neuronal development, and neuroprotection and is a well-studied protein family in AD CSF^31,32,33^. Our longitudinal analysis shows that altered plasma expression of the 14-3-3 protein family is an age-related change, being observed in all three longitudinal analysis groups (cognitively normal controls, MCI and AD), so it may have functional implications for progression to MCI and/or AD since ageing is the major risk factor for these conditions. However, as the hippo family members are not unique to AD (Table S3, Figure S5), they are less likely to be valuable biomarkers.

Damaged and misfolded proteins accumulate during the ageing process, affecting cell function and tissue homoeostasis. Cellular clearance processes such as the proteasome are a critical component of the proteostasis network, involved in the degradation and recycling of damaged proteins. Proteasome activity declines with age, and dysfunctional proteasomes are related to late-onset diseases^34^. We identified five dysregulated proteasome members, all of which were upregulated in plasma: PSMA4, PSME2, PSMA2, PSMA5, and PSMB8, suggesting that intracellular protein turnover is compromised with ageing. Other protein families of which multiple members were identified in our longitudinal ageing groups include actin-related protein (6 DEPs), chloride intracellular channel protein (2 DEPs), glutathione S-transferase (2 DEPs), L-lactate dehydrogenase (2 DEPs), peptidyl-prolyl cis-trans isomerase (2 DEPs), protein S100 (2 DEPs), and rho GDP-dissociation inhibitor (2 DEPs) (Table 2 and Table S3). Since most of these are intracellular proteins, their presence in plasma is likely a reflection of cellular senescence, increased fragility and cell death with ageing. Therefore, the functions they sub-serve are likely compromised with ageing and may predispose to disease progression. However, since they were also present in the cognitively normal ageing group, these changes are insufficient on their own to explain progression to cognitive disorder or neurodegenerative disease. Changes to many of these proteins have previously been attributed to associations with either MCI and/or AD^35–37^. The current work demonstrates the need for particular care in age matching in case-control studies, especially biomarker studies.

A variety of other age-related protein changes were observed in common across all three longitudinal analysis groups (normal controls, MCI and AD), including the HIF-1 signalling pathway (Table S4) and several proteins abundant in the CNS (NCAM1, YWHA family, PKM, NME2, MAPRE1) indicating that age-related changes within the CNS can be detected in plasma, with techniques which allow sufficient depth of proteome coverage. Previously, studies showed that HIF-1 generates a deficit in mitochondrial biogenesis during the ageing process impairing energy-dependent cellular functions such as cell and tissue repair^38^. We have identified a list of markers such as TIMP1, GAPDH, ENO1, PGK1, LDHB, and LDHA that can aid in the understanding of mitochondrial dysregulation in ageing (Table S4). In addition, a HIF-1 signalling pathway is known to both promote and limit longevity via pathways that are mechanistically distinct using hypoxic response transcription factor HIF-1^39,40^. Validation of these DEPs in large sample size cohorts might improve our understanding of human ageing. Such broad-ranging pathway changes in ageing may also help explain why ageing is the single major risk factor for a wide variety of diseases, including age-related neurodegenerative diseases such as dementia. This wide range of pathways impacted by the ageing process likely overlaps with many disease processes, making ageing an accelerant if not a causative risk factor for cognitive decline and/or disease.

### Changes in the plasma proteome associated with progression to incipient MCI and AD dementia over 6 years

To identify MCI and AD specific changes in the longitudinal analysis groups, we filtered the DEPs list for age-independent protein changes, which were unique to either AD or MCI, but not similarly changed in cognitively normal ageing (Figure 2, Tables S5a and S6a, and Table 2). A characteristic of AD patients that brain imaging techniques can detect is impaired glucose uptake in brain regions with neuritic plaques^41–43^. Numerous AD-specific DEPs involved in metabolism (e.g., APOD, ALDOB, MAN2A2, GPX3, HPRT1, ALDH1A1, AMY1A, MGAT1, and IGFBP5) were elevated in plasma in our investigation, reflecting impairment of the cellular metabolic processes in which these proteins partake (Table S7).

APOD is a glial-expressed lipid transport protein of the lipocalin family that has been shown to protect against oxidative stress^44^. Our longitudinal data show that increased APOD is observed in MCI and AD plasma, but not in cognitively normal ageing (Tables S5a and S6a). This observation is consistent with other published work, which shows elevated APOD in AD, Parkinson’s disease, Schizophrenia, Stroke and Bipolar disorder^44,45,46^. Increased plasma IGFBP5 also appears to be related to cognitive decline since older adults with depression lose cognitive abilities faster when they have higher IGFBP-5 levels^47^. HPRT1 was recently identified as one of the most strongly validated metabolic proteins, exhibiting a substantial increase in AD CSF cohorts, demonstrating a direct link between energy production and synaptic signalling at the neuronal membrane^32,48^. Another metabolic protein identified is ALDH1A1, a multifunctional enzyme with dehydrogenase, esterase, and antioxidant activities and critical for normal brain homeostasis, which was upregulated in AD downregulated in MCI in our data. A recent study shows neurons may upregulate ALDH1A1 activity to compensate for oxidative stress-induced damage in the brain^49^. We are proposing metabolic abnormalities, which can be identified in plasma, as a critical component of longitudinal AD aetiology, a better understanding of which might provide novel metabolic targets for therapeutic development.

In addition to metabolism, a large group of proteins were associated with homeostasis in AD and MCI (Table S7 and S8). Homeostasis related proteins were upregulated in MCI but downregulated in AD. A recent study suggested that firing instability and poor synaptic plasticity during the early stages of AD initiate a vicious loop that results in integrated homeostatic network (IHN) dysregulation^50^. According to this idea, the collapse of the IHN is the primary factor driving the transition from early memory deficits to neurodegeneration^51^. Homeostatic proteins which were downregulated in AD are COL1A1, SELL, ENDOD1, TF, SERPINA1. Decreased level of TF in AD plasma and brain samples has been reported previously^52,53^. Consistent with these early reports, we found significant downregulation of TF during longitudinal progression to AD in our study.

In MCI, several unique homeostasis markers were differentially expressed, including upregulation of RAB27B and PPBP, which may participate in the pathology underlying MCI. RAB27A and RAB27B are involved in the docking of MVE at the plasma membrane^54^. Previous studies have found that the upregulated expression of RAB27 correlated with antemortem indicators of cognitive deterioration in MCI and AD brains^54–56^. In our data, RAB27B was elevated in MCI but did not change in AD, which may imply endosomal dysfunction as an early change detected in MCI, which may contribute to progression to AD in later stages of life. Alternatively, it may also be a change specific to MCI, and studies of longer duration may help decipher what changes are associated with stable MCI vs progression to AD.

It is believed that the extracellular matrix contains collagens, which are essential in axonal guidance, synaptogenesis, cell adhesion, the formation of brain architecture and neural maturation^57–59^. A gene from the college family, *COL25A1,* was overexpressed in neurons of transgenic mice leading to AD-like brain pathology^60^. In our data, COL1A1 was upregulated in longitudinal MCI progression but downregulated in AD. Such differences may point to mechanisms that help limit the level of impairment and avoid progression to greater levels of cognitive impairment.

Moreover, SERPINA1 is emerging as a key neuroinflammation modulator^61^, also reported being released from the brain tissue to the CSF^62^. Higher CSF levels of SERPINA1 have been linked to the clinical diagnosis of AD^63^. Here we found that SERPINA1 was upregulated in MCI and also in preclinical AD but downregulated in clinical AD (Figure 5B), suggesting that it may be an early marker of synaptic loss particularly evident in plasma at preclinical stages of dementia, at a time when much damage is in active progress, and plateauing/declining in parallel with the onset of clinical symptoms.

### Proteomic changes in clinically diagnosed AD and MCI relative to their age-matched cognitively normal controls (potential clinical diagnostic markers)

A total of 70 and 89 DEPs were identified in a cross-sectional analysis of incipient AD (ADW4/CRTLW4) and incipient MCI (MCIW4/CRTLW4), respectively, indicating that a potentially rich biomarker signature for AD and MCI is available in plasma.

Of the 70 cross-sectional AD-associated DEPs, 15 were also identified in longitudinal AD analysis (Figure S6), of which 11 had a consistent direction of expression change. These may be the most robust biomarker candidates as they change consistently in longitudinal and cross-sectional AD relative to their age-matched controls (Figure S6 and Table S5c). Of these, 10 DEPs (TF, VCP, PSMB2, PA2G4, PAM, MAN2A2, TNXB, FAM3C, ALDOB, and QDPR) were enriched with extracellular exosome GO terms in the STRING analysis. At an early stage of AD, a rise in the protein levels of total and phosphorylated tau in exosomes has been found in the CSF^64,65^. Another finding implies that exosomes may be the primary mechanism controlling the spread of Aß and tau^66^. Our findings are consistent with the published literature, which indicates that exosome dysregulation is a key event in AD patients compared to their age-matched healthy controls^65^. In addition to homeostasis and metabolic disruption, neutrophil degranulation, protein binding, and transport were the most enriched pathways in incipient AD-related DEPs in crosssectional analysis. Neutrophil activation and accompanying oxidative stress have been linked to AD pathogenesis^67^. It is noteworthy that our study identified brain-derived proteins such as MAN2A2, PAM, TF, QDPR, FAM3C, which have previously been reported to be dysregulated in AD CSF and brain^68^.

There were 18 DEPs common to both MCI and AD, which may be considered potential shared biomarkers of cognitive change, including FAM3C, TF, CLIC1, CHI3L1 PRDX4 and others (Figure 3D and E), and possibly reflecting the underlying disease process. FAM3C is an interleukin-like protein (also called ILEI) with a proposed role as a metabolic regulator^69^. FAM3C ameliorates Aß pathology by reducing Aß levels^70^, has been suggested as a surrogate biomarker of Aβ in the brain^71^ and FAM3C levels are lower in the AD brain^72^. Its normal expression level is exceptionally high in the gut, thyroid and brain (https://www.ncbi.nlm.nih.gov/gene/10447). Previous work has reported lower levels of CSF FAM3C in AD and MCI groups which suggested this may result in a build-up of Aβ in the brain and eventual development of AD^72^. We observed a higher level of FAM3C in plasma samples of MCI and AD compared to respective age-matched controls suggests either loss from the CNS or a homeostatic/compensatory increase in response to loss in an organ system/s.

The protein CHI3L1 (also known as YKL-40) is secreted by activated microglia and reactive astrocytes^73^ and is thought to have a role in inflammation and tissue remodelling, particularly angiogenesis^74^. In the current work, CHI3L1 was increased in both MCI and AD plasma, consistent with reported observations of higher CHI3L1 levels in AD than in healthy controls or MCI patients^75^, and also of other neurodegenerative diseases^76^. In addition, several studies have reported that CHI3L1, an astrocyte-derived protein, is increased in AD CSF and has been suggested to be a marker for progression from MCI to AD^77,78^.

Serotransferrin (TF) decreased in incipient AD while increased in MCI^53^. Transferrin (Tf) is an important iron-binding protein that is thought to have a critical function in iron ion (Fe) absorption via the transferrin receptor (TfR). Elevated Fe levels in AD brains have been reported and linked to amyloid plaque formation^79^.

### Potential early disease markers; preclinical changes in AD

Preclinical AD, defined as a stage of neurodegeneration occurring before the onset of clinical symptoms, is likely to be the more effective time point to apply potentially disease-modifying interventions in AD^80,81^. A total of 160 DEPs of preclinical AD (ADW1/CTRLW1) were identified (Figure 5B), which were a considerably larger number than the 71 DEPs identified in incipient AD (Figure 4A). The considerably larger pool of preclinical AD-associated DEPs may in part be evidence of pathology in progress, in addition to providing several putative early biomarkers. Reasoning that the most robust biomarkers may be those that continue to be observed with clinical disease onset, 15 DEPs were shared with clinical AD (ADW4/CTRLW4) (Figure 5C, 5D and Table S12b). Furthermore, of these 15 DEPs, 8 were unique to preclinical and clinical AD but not identified in clinical MCI (MCIW4/CTRLW4). These 8 proteins were PGK1, ALDOB, VCL, PAM, PSMB2, VCP, MAPRE2, and ISOC1, and may be specific to AD-related pathology, rather than just associated with cognitive decline *per se.* Interestingly, glycolysis and gluconeogenesis presented as top GO terms with significant enrichment in preclinical and clinical AD plasma. This concurs with the presence of three glycolytic proteins from our 8-protein signature: PGK1, VCP and ALDOB. Numerous studies have demonstrated dysregulation of glucose metabolism in the brain, which has long been recognised as an apparent anomaly that commences during the preclinical stage of AD^82,83,84^ and remains a feature with incipient AD. Apart from the well-known CSF AD biomarker (CHI3L1), we propose a list of novel markers, including PSMB2, PAM, ALDOB, TF, MAPRE2, VCP, which may be potential preclinical biomarkers for the identification of AD, being dysregulated in all three AD comparison groups, i.e., longitudinal (ADW4/ADW1), incipient AD (ADW4/CTRLW4) and preclinical AD (ADW1/CTRLW1). That they are all representative of different aspects of AD pathology (PSMB2, proteasomal turnover of dysfunctional proteins; PAM, signalling peptide synthesis; ALDOB, glycolysis and gluconeogenesis metabolic pathways; TF, iron-binding and transport; MAPRE2, a microtubule-associated protein with a possible role in signal transduction; VCP, segregation of proteins for degradation by the proteasome) may offer a specificity advantage, since AD is a complex multifactorial disease with dysfunction of multiple cellular pathways. It is of note that of these 6 proteins, the only DEP with a consistent fold-change (increased) across all three groups is PSMB2.

### Mechanisms of AD

MCI is often considered a risk factor and/or prodromal stage of AD^85^, so it was of note that in the comparisons of AD and MCI, only approximately 18% (19 proteins) and 13% (18 proteins) of DEPs were common to both AD and MCI in the longitudinal (Figure 3A) and cross-sectional analyses (Figure 3D). In this context, it is of interest that by far most DEPs identified in AD and MCI are specific to each condition rather than shared.

Another prevalent hypothesis is that dysfunction of the cytoskeleton and microtubule system may contribute to AD pathogenesis^86,87^. MAPRE2, a microtubule-associated protein, and VCL, a membrane-cytoskeletal protein, are involved in microtubule polymerization, cell-cell and cellmatrix junctions. A recent study suggested that MAPRE2 is involved in cellular migration of cranial neural crest cells, among others, via its involvement in focal adhesion dynamics^88^, although no direct association between MAPRE2 and AD progression has been established previously.

Dysregulation of phosphorylation in AD is commonly observed^89^, so it is interesting that several proteins in our preclinical biomarker list are directly or indirectly involved in phosphorylation (PGK1, GANAB, PPP2R4). Several studies have reported that GANAB and PPP2R4 are dysregulated in AD CSF and brain^90,91^. Most of the brain Ser/Thr phosphatase activity involves PP2A family enzymes. The dysfunction of PPP2R4 has been linked to tau hyperphosphorylation, amyloidogenesis and synaptic deficits that are pathological hallmarks of AD^91,92^. Furthermore, SPARC and ARPC5 are proteins involved in regulating cell-cell interactions, actin polymerisation and neural plasticity, respectively. It has been reported that chronic stress significantly increased the level of ARPC5 in the hippocampus, implying that chronic stress-induced alterations in hippocampal proteins are related to synaptic plasticity^93^.

### Picking candidate biomarkers specific to incipient MCI and AD and preclinical AD

Proteomic expression change was seen in a surprisingly large number of plasma proteins in normal ageing over the 6-year period of this study (71 DEPs, Table 2). Even after these were excluded from the MCI and AD longitudinal analyses, we were still left with a long lists of proteins (66 and 60 MCI and AD specific DEPs, respectively), and similarly large numbers in the cross-sectional analyses (89 and 70 MCI and AD specific DEPs respectively). Such abundance presents a dilemma as to which DEPs might be ideal biomarker candidates. One approach is to select potential candidates based on consistency of fold-change in both longitudinal and cross-sectional analyses for each of the incipient AD and MCI and preclinical AD groups. These are much shorter lists but are likely much stronger candidates for future validation work. It is of note that most DEPs in the incipient MCI and AD groups, are upregulated (~ 66% each). In contrast, DEPs with consistent fold changes in the preclinical and incipient AD groups are mostly downregulated, with only 2 DEPS (< 20%) upregulated, these two being the cell-matrix protein CH13L1 and the proteasome protein PSMB2.

#### Limitations

The study’s limitations included 1). The depth of the approach restricted the number of samples that could be studied, reducing the power of the analysis. 2). MCI and AD were diagnosed by consensus and met the NIA-AA criteria for MCI and AD, respectively; however, no biomarker confirmation was available. 3). The results were not replicated in independent cohorts. As this is an exploratory study, additional research into the relevance of these proteins is warranted in prospective studies of dementia-free individuals in midlife and long-term dementia incidence follow-up.

#### Conclusion

We have identified an in-depth plasma proteome showing we can now detect proteins generated from the brain and generate protein signatures. These protein changes are consistent across different independent search engines, paving the path for future research on neurodegenerative disorders biomarker identification. Many studies showing that CSF proteome can more closely reflect the brain disease pathology, however, the relationship of plasma proteome to brain changes is still insufficiently understood. Global deep plasma proteomics analysis, in combination with longitudinal and cross-sectional analyses of an older age cohorts, ageing with normal cognition or progressing to MCI or dementia, revealed changes common to ageing regardless of diagnosis, and molecular similarities and differences AD and MCI, as well as some putative dementia specific plasma biomarkers for clinical and preclinical AD. The considerably larger pool of preclinical AD associated differentially expressed proteins may in part be evidence of pathology in progress, in addition to providing a large pool of putative early biomarkers. Apart from the well know CSF AD biomarker (CHI3L1) we propose a list of novel markers, including PSMB2, PAM, ALDOB, TF, MAPRE2 which may be potential AD preclinical biomarkers for the identification of AD dementia.

## Supporting information

Venn diagrams showed an overlap of 1,467 proteins in the three longitudinal comparisons

Scatter plots and density plots for all 6 comparisons are shown in Figures S2i and S2ii respectively

Scatter plots and density plots for all 6 comparisons are shown in Figures S2i and S2ii respectively

The detailed scatter plots were plotted using the DEPs from both the search engines i.e. PD2.4 and scaffold in all 6 comparisons Figure S2iii

These 71 age-related DEPs were manually grouped into 12 protein functional categories based on gene ontology (GO) using the PD2.4 analysis outcomes (F

followed by colloidal coomassie G250 staining25 (Figure S4).

Of 71 DEPs in normal ageing, only two proteins were decreased, these being methanethiol oxidase (SELENBP1) and neuronal adhesion molecule 1 (NCAM1) Fi

Relatively few of the same DEPs were identified in both the cross-sectional and longitudinal analyses of AD and MCI, being 15 (Figure S6A and S6B) and

we identified 450 differentially expressed proteins (DEPs) in the longitudinal comparisons and 553 DEPs in the cross-sectional comparisons, summarized

## Author Contributions

A.P. and P.S. conceived the project; G.K. and A.P. designed the experiments; G.K. performed the chromatographic and SDS/PAGE fractionation experiments and preparation of peptides for LC-MS/MS; A.P. acquired the LC-MS/MS; G.K. and A.P. conducted bioinformatics (PD2.4 and Scaffold) data analysis; G.K., A.P. wrote and revised the manuscript; and P.S. supervised the project and reviewed the manuscript. All authors read, edited, and revised the manuscript and accepted the final version.

## Acknowledgements

G.K. is thankful to the UNSW Scientia fellowship. We would also like to thank the Bioanalytical Mass Spectrometry Facility, University of New South Wales, for providing the necessary facilities to carry out the research work. G.K. is thankful to Jitender Singh Virk and Syed Azmal Ali for their help in learning R studio and programming language for making figures.

## Supplementary Figures

**Figure S1:**

**A.** Box and Wisker plots of abundance values of all 66 individual samples. The line within the box denotes the median value, and the upper and lower ranges of the box indicate the 5 and 95 percentiles of the abundance values, respectively (output from ProteomeDiscoverer 2.4 software).

**B.** The similarity matrix and heat map were constructed using the Pearson correlation values of the 7 comparisons, clustered based on the *k* means algorithm.

**C.** Venn diagrams depicting the total number of proteins identified in longitudinal comparison. We identified 1467 proteins which were common in all three longitudinal groups i.e., normal ageing, MCI and AD.

**D.** Venn diagrams depicting the total number of proteins identified in cross-sectional comparisons. We identified 903 proteins which were common in all four cross sectional groups i.e., preclinical AD, MCI, AD and MCI+AD.

**Figure S2i:** Scatter plot and regression analysis of abundance ratio of DEPs in both PD2.4 and scaffold; CTRLW4/CTRLW1, MCIW4/MCIW1, ADW4/ADW1, MCIW4/CTRLW4, ADW1/CTRLW1, and ADW4/CTRLW4.

**S2ii:** Density plot and regression analysis was plotted between all 6 comparisons of normal control, MCI, and AD in both longitudinal and cross-sectional comparisons. Each dot represents abundance ratio of each protein, and the colour shows the dot density. **S2iii.** Scatter plots were plotted using only DEPs from each comparison. **A.** 71 DEPs from CTELW4/CTRLE1 (Table S3) **B.** 66 DEPs from MCIW4/MCIW1 (Table S6a) **C.** 60 DEPs from ADW4/ADW1 (Table S5a), **D.** 89 DEPs from MCIW4/CTRLW4 DEPs (Table S6b), **E.** 70 DEPs from ADW4/CTRLW4 (Table S5b), **F.** 160 DEPs from ADW1/CTRLW1 (Table 12a)

**Figure S3:** The final list of DEPs from longitudinal comparisons were sorted into lists based on the protein function classes i.e., extracellular function, cell adhesion, growth factors, cell signaling, neuroinflammation, cytoskeleton, protein turnover, DNA binding repair, metabolism, membrane trafficking, neuron and synapse, antioxidant activity: **A.** DEPs specific to longitudinal ageing, and common to all three longitudinal clinical groups, including ageing over the 6 year time period between W1 and W4 with stable normal cognition and progression to MCI or AD **B.** DEPs unique to longitudinal progression to MCI. These DEPs are not found in the normal ageing or AD groups **C.** DEPs unique to longitudinal progression to AD. These DEPs are not found in the normal ageing ot MCI groups.

**Figure S4:** Representative image of NuPAGE LDS gel profile of and depleted plasma containing low abundant plasma proteins (LAP) from HU14 from all the individuals. Each gel lane contained and equal loading of total protein (50 ug total proteins were loaded per gel lane).

**Figure S5: A.** Heatmap of 71 dysregulated proteins containing 69 upregulated and 2 downregulated in similar direction in all normal ageing, MCI and AD showing the plasma proteome changes with age and not specific to disease. **B:** Volcano plots highlight the 20 DEPs with highest fold change in longitudinal ageing (we have highlighted only top 20 proteins to avoid the overcrowding on volcano plots) full list of DEPs with age are presented in Table S3. **C:** Upregulated GO enrichment of pathways linked to ageing are presented in this figure, however only 2 DEPs were downregulated in ageing, no GO enrichment was identified in STRING software for downregulated proteins. The full list of GO enrichment is presented in Table S4.

**Figure S6: A.** Venn diagram showing the number of common DEPs in longitudinal and cross sectional AD. **B.** The list showing the common 15 DEPs with gene name and fold change in both longitudinal and cross sectional AD. **C.** Venn diagram showing the number of common DEPs in longitudinal and cross sectional MCI. **D.** The list showing the common 16 DEPs with gene name and fold change in both longitudinal and cross sectional MCI.

**Table S1:** Total number of proteins identified in PD2.4 and scaffold search engines in all 7 comparisons.

**Table S2:** List of DEPs those quantified in >6 individuals, proteins identified with a minimum of two peptides/protein, consistent direction of protein fold change across two bioinformatics platforms with orthogonal quantification approaches (peak area ratio with PD2.4 and spectral counting with Scaffold) with a fold change of at least 20% (≤0.08 and ≥ 1.2) in preferably both search engines or at least one. **S2a.** Using this approach, we identified 450 differentially expressed proteins (DEPs) in longitudinal comparisons, **S2b.** 553 DEPs in cross-sectional comparisons, **S2c.** 297 common between both comparisons.

**Table S3:** The final list of longitudinal 71 DEPs in normal ageing (changed in similar direction in longitudinal normal ageing, MCI and AD were considered as ageing related changes, not specific to the disease).

**Table S4:** GO enrichment was performed using 71 DEPs in normal ageing to gain the insights into the biological processes, cellular components, molecular functions, KEGG and Reactome molecular pathways affected in normal ageing using STRING software.

**Table S5a:** The list of proteins uniquely differentially expressed in AD in both longitudinal (ADW4/ADW1) and **S5b.** cross-sectional comparison (ADW4/CTRLW4) and **S5c.** DEPs which are common in both AD comparisons.

**Table S6:** List of differentially expressed proteins (DEPs) in ageing at W1 with progression to MCI at W4. The proteins in these tables were quantified in >6 individuals, proteins identified with a minimum of two peptides per protein, consistent direction of protein fold change across two bioinformatics platforms with orthogonal quantification approaches (peak area ratio with PD2.4 and spectral counting with Scaffold) with a fold change of at least 20% (≤0.08 and ≥ 1.2) in preferably both search engines or at least one. **S6a:** The list of proteins uniquely differentially expressed in MCI in both longitudinal (MCIW4/MCIW1) and **S6b.** cross-sectional comparison (MCIW4/CTRLW4) and **S6c.** the list of DEPs which are common in both MCI comparisons.

**Table S7:** GO term enrichment was performed using 60 DEPs identified in the longitudinal AD group (Table S5a), 39 of which were upregulated and 21 of which were downregulated. The GO term enrichment approach, was applied to disease associated DEPs to gain insight into biological processes, cellular components, molecular functions, KEGG and Reactome molecular pathways associated with longitudinal AD. STRING software was used for GO term enrichment analysis.

**Table S8:** GO enrichment was performed using the 66 proteins which were differentially expressed in MCI (W4/W1, longitudinal analysis), with 41 upregulated and 25 downregulated to gain the insights into the biological processes, cellular components, molecular functions, KEGG and Reactome molecular pathways affected in longitudinal MCI using STRING software. The protein list used for the enrichment analyses in this table is shown in Table S6a.

**Table S9:** The list of proteins differentially expressed in both longitudinal and cross sectional AD and MCI analyses. Proteins listed here are quantified in >6 individuals, identified with a minimum of two peptides per protein, consistent direction of protein fold change across two bioinformatics platforms with orthogonal quantification approaches (peak area ratio with PD2.4 and spectral counting with Scaffold) with a fold change of at least 20% (≤0.08 and ≥ 1.2) in preferably both search engines or at least one. **S9a.** A total of 19 DEPs were common in both the longitudinal AD and MCI plasma proteome profiles. **S8b.** The list of 18 DEPs were common in both the cross sectional AD and MCI.

**Table S10:** GO enrichment was performed using 70 DEPs identified in cross sectional analysis of incipient AD (ADW4/CTRLW4) including 27 upregulated and 43 downregulated to gain the insights into the biological processes, cellular components, molecular functions, KEGG and Reactome molecular pathways associated with incipient AD. Table S5b shows the DEPs used in the enrichment analyses presented here.

**Table S11:** GO enrichment was performed using 89 proteins were differentially expressed in cross sectional MCI, with 53 upregulated and 36 downregulated (Table S6b) to gain the insights into the biological processes, cellular components, molecular functions, KEGG and Raectome molecular pathways affected in longitudinal cross sectional MCI using STRING software.

**Table S12:** The list of proteins differentially expressed in preclinical AD (ADW1/CTRLW1), which is a comparison of the AD baseline data, at which time point subjects were cognitively normal, with aged matched controls (W1 baseline values for individuals who remain cognitively stable over the 6 year period of the study). This table list differentially expressed proteins (DEPs) quantified in >6 individuals, proteins identified with a minimum of two peptides per protein, consistent direction of protein fold change across two bioinformatics platforms with orthogonal quantification approaches (peak area ratio with PD2.4 and spectral counting with Scaffold) with a fold change of at least 20% (≤0.08 and ≥ 1.2) in preferably both search engines. **S12a:** In preclinical AD, a total 160 were dysregulated proteins including 59 upregulated and 101 downregulated proteins in preclinical AD (ADW1/CTRLW1). **S12b**. A total of 15 that were found to be commonly differentially expressed in age-matched clinical (cross sectional AD) and preclinical AD protein lists. **S12c**. A total of 16 DEPs were identified as frequently dysregulated in preclinical AD and cross sectional MCI.

**Table S13:** In preclinical AD (W1AD/W1CTRL), GO enrichment was performed using 160 dysregulated proteins, including 59 upregulated and 101 downregulated proteins listed in Table S12a, to gain the insight into the biological processes, cellular components, molecular functions, KEGG and Reactome molecular pathways associated with preclinical AD. GO term enrichment was performed using STRING software.

