## Supplementary figures and images for "Deep proteome analysis of plasma reveals novel biomarkers of mild cognitive impairment and Alzheimer’s disease: A longitudinal study"

### followed by colloidal coomassie G250 staining25 (Figure S4).

#### Figure S4

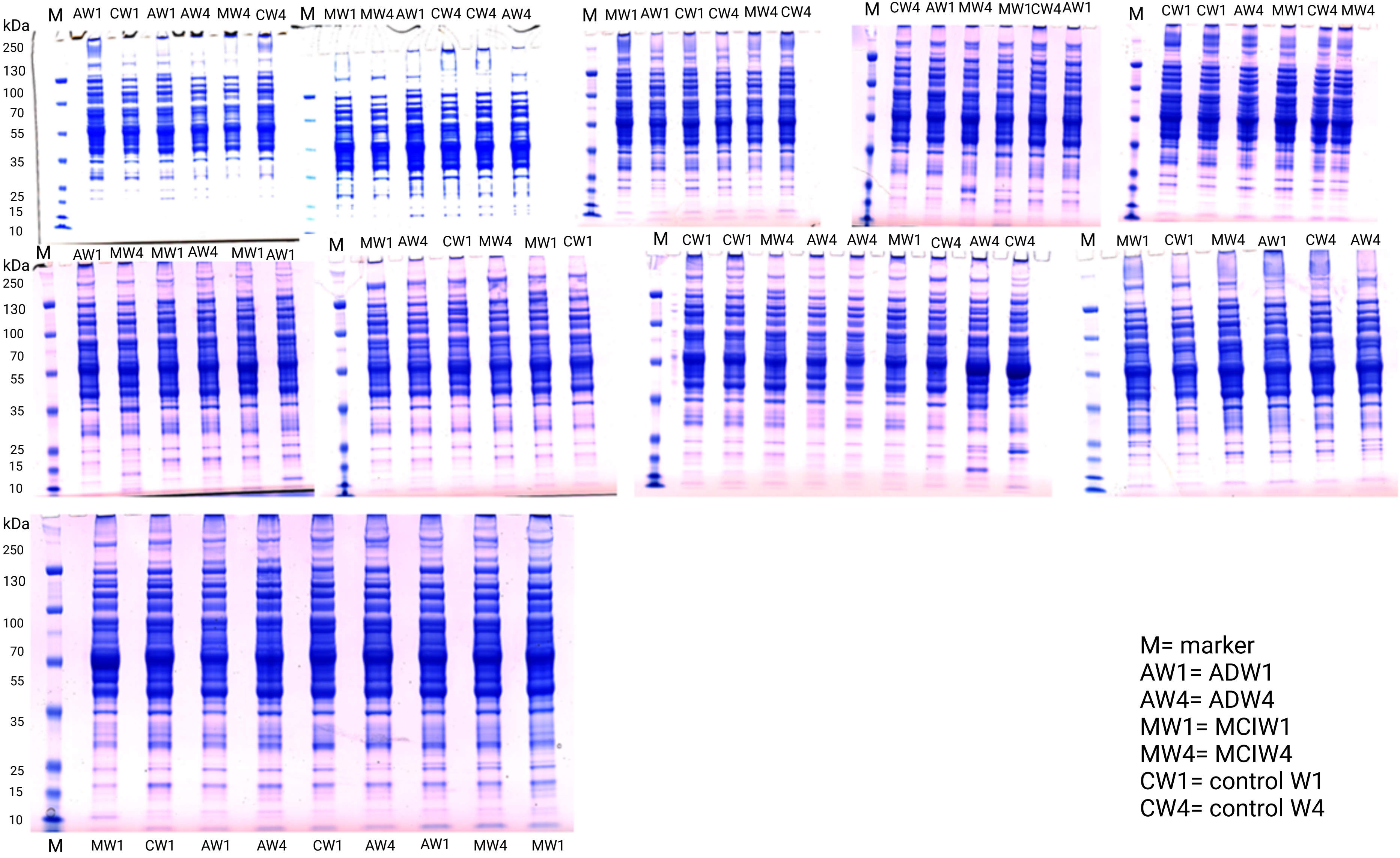


M= marker
AW1= ADW1
AW4= ADW4
MW1= MCIW1
MW4= MCIW4
CW1= control W1
CW4= control W4

### Of 71 DEPs in normal ageing, only two proteins were decreased, these being methanethiol oxidase (SELENBP1) and neuronal adhesion molecule 1 (NCAM1) Fi

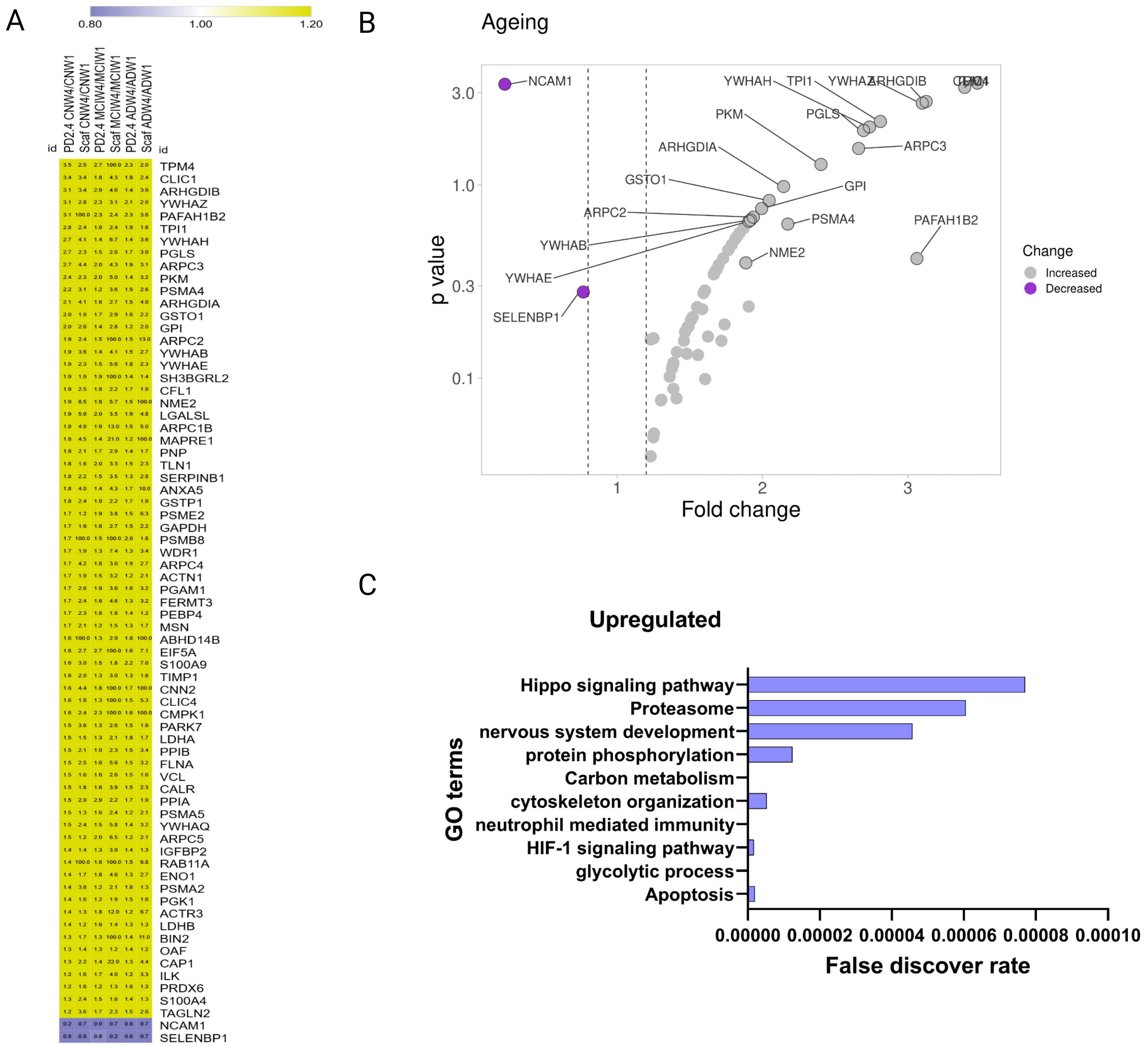

### Scatter plots and density plots for all 6 comparisons are shown in Figures S2i and S2ii respectively

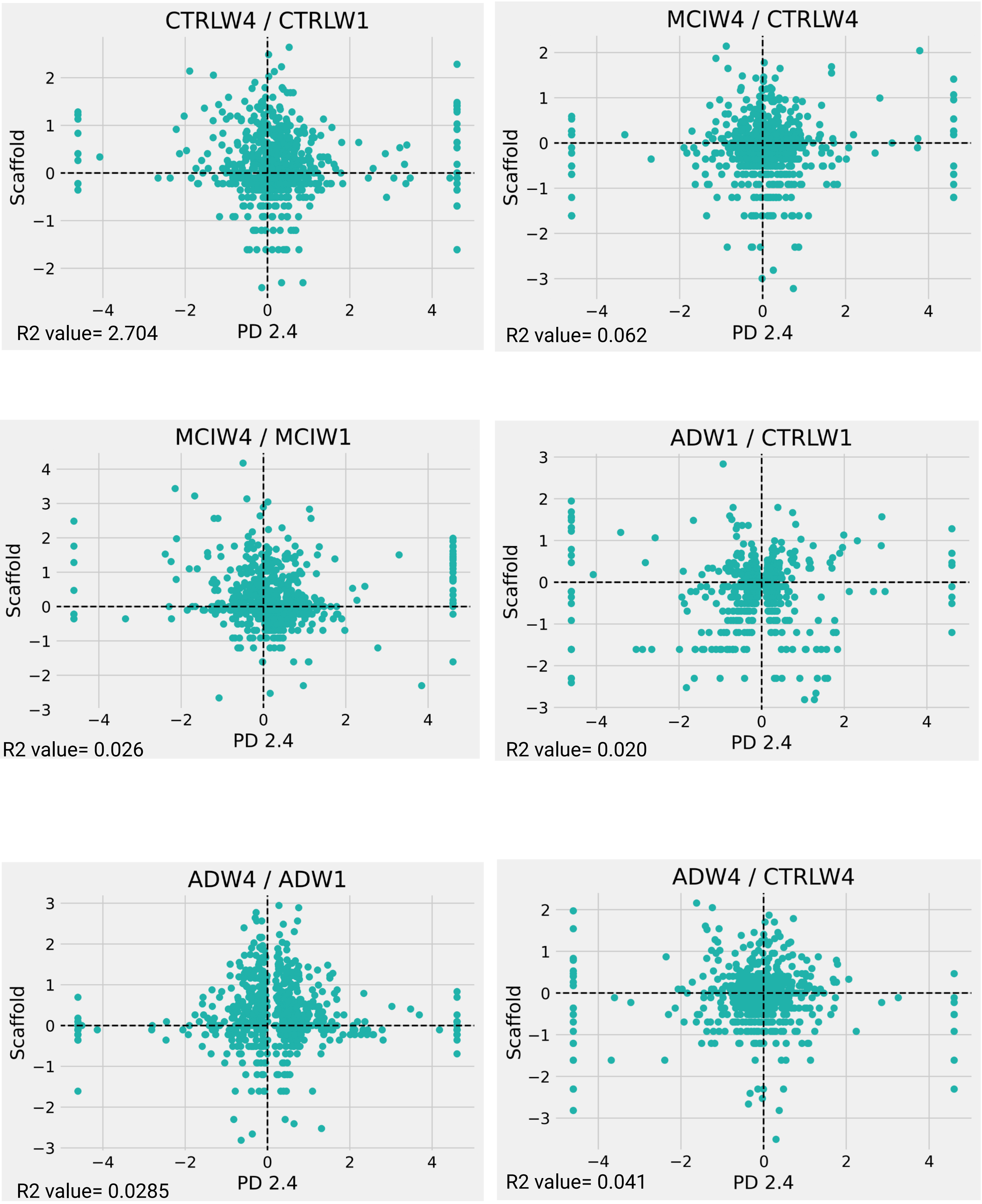

### Scatter plots and density plots for all 6 comparisons are shown in Figures S2i and S2ii respectively

Figure S2ii

R2 value= 0.053

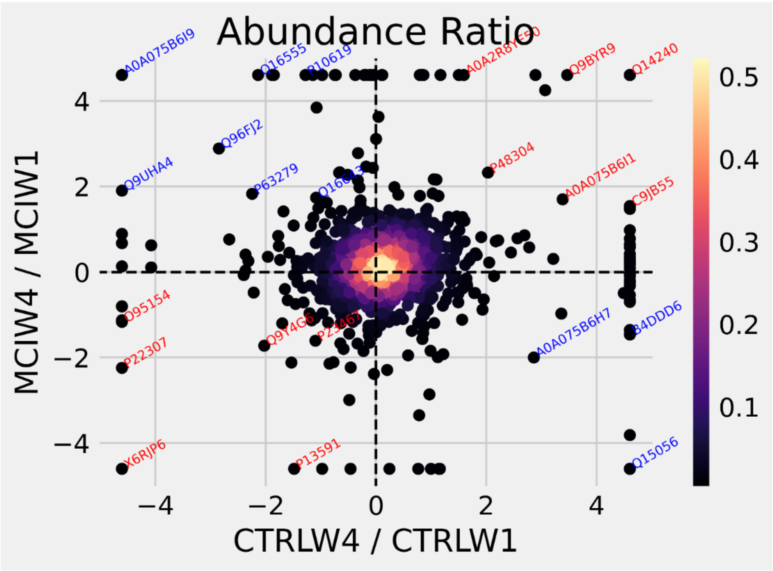


R2 value= 0.125

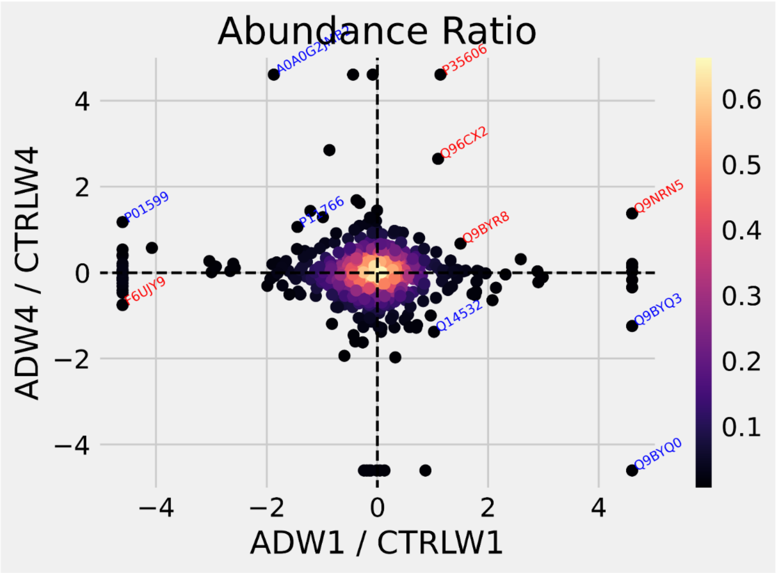


R2 value= 0.011

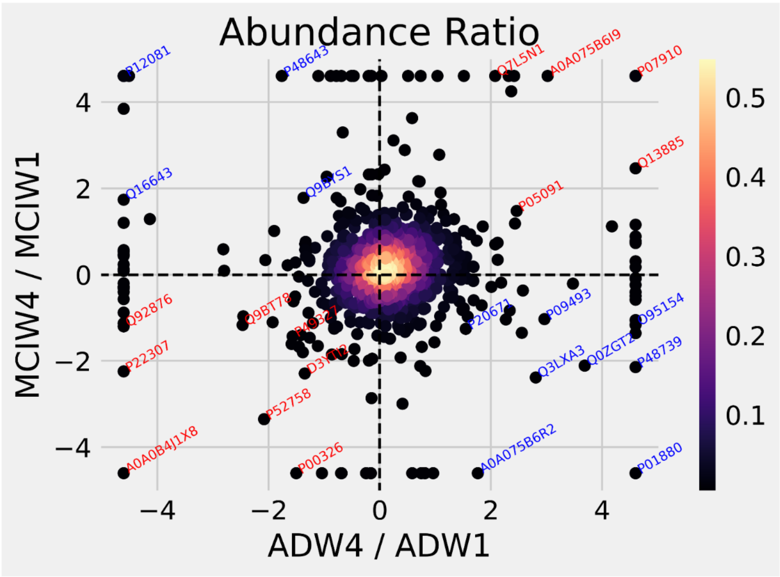


R2 value= 0.056

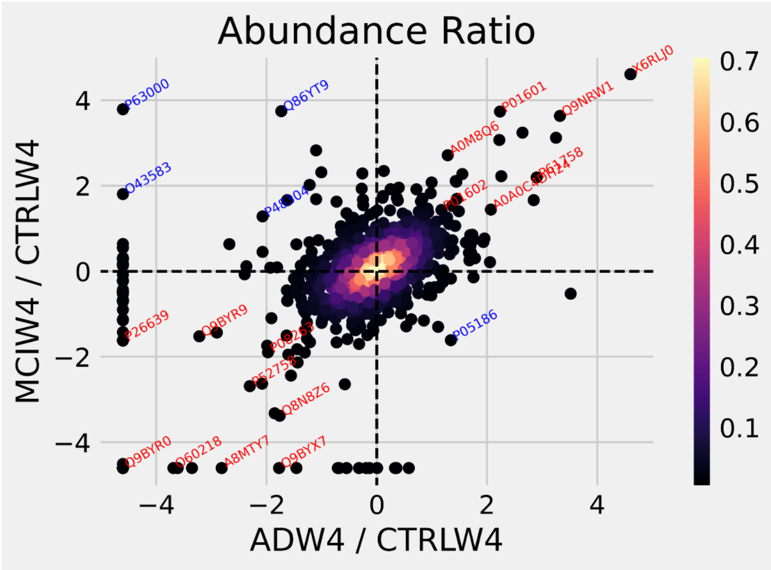


R2 value= 0.0249

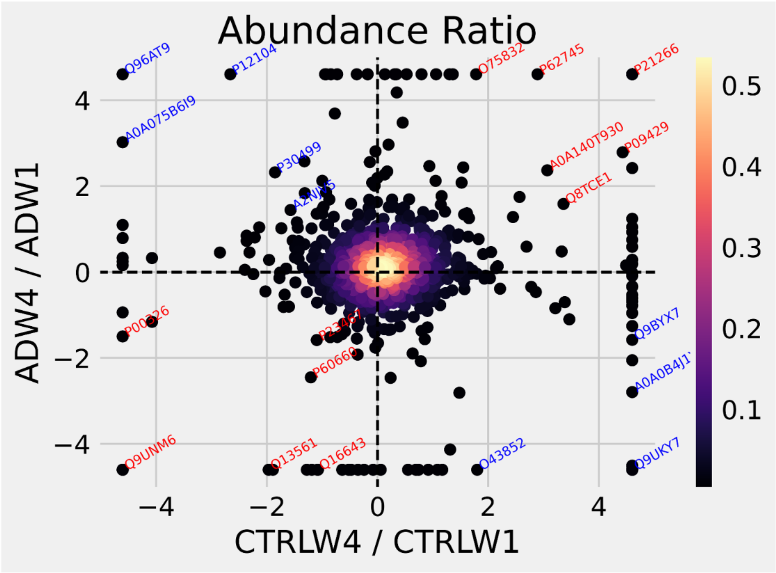

### The detailed scatter plots were plotted using the DEPs from both the search engines i.e. PD2.4 and scaffold in all 6 comparisons Figure S2iii

Figure S2iii

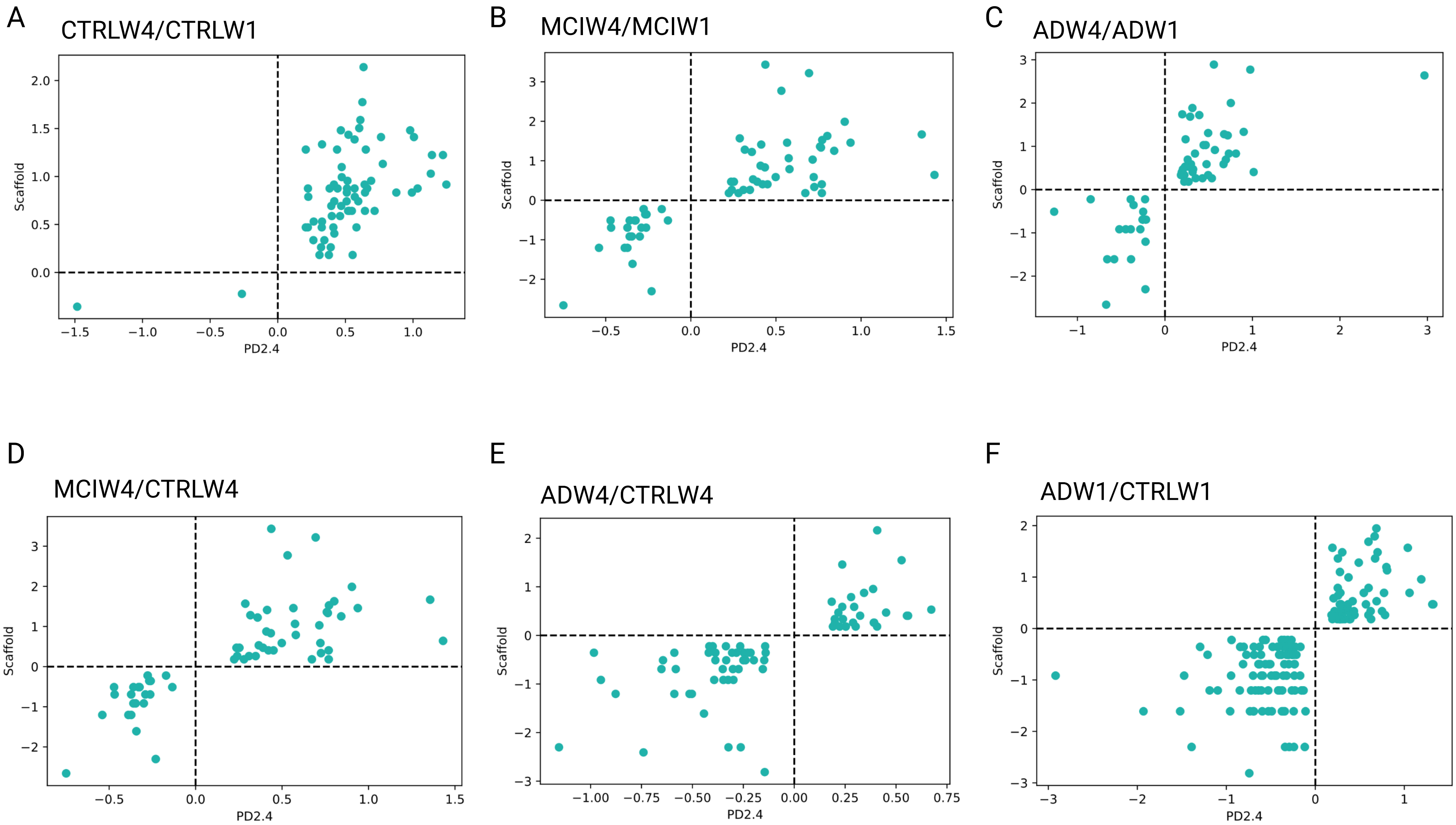

### These 71 age-related DEPs were manually grouped into 12 protein functional categories based on gene ontology (GO) using the PD2.4 analysis outcomes (F

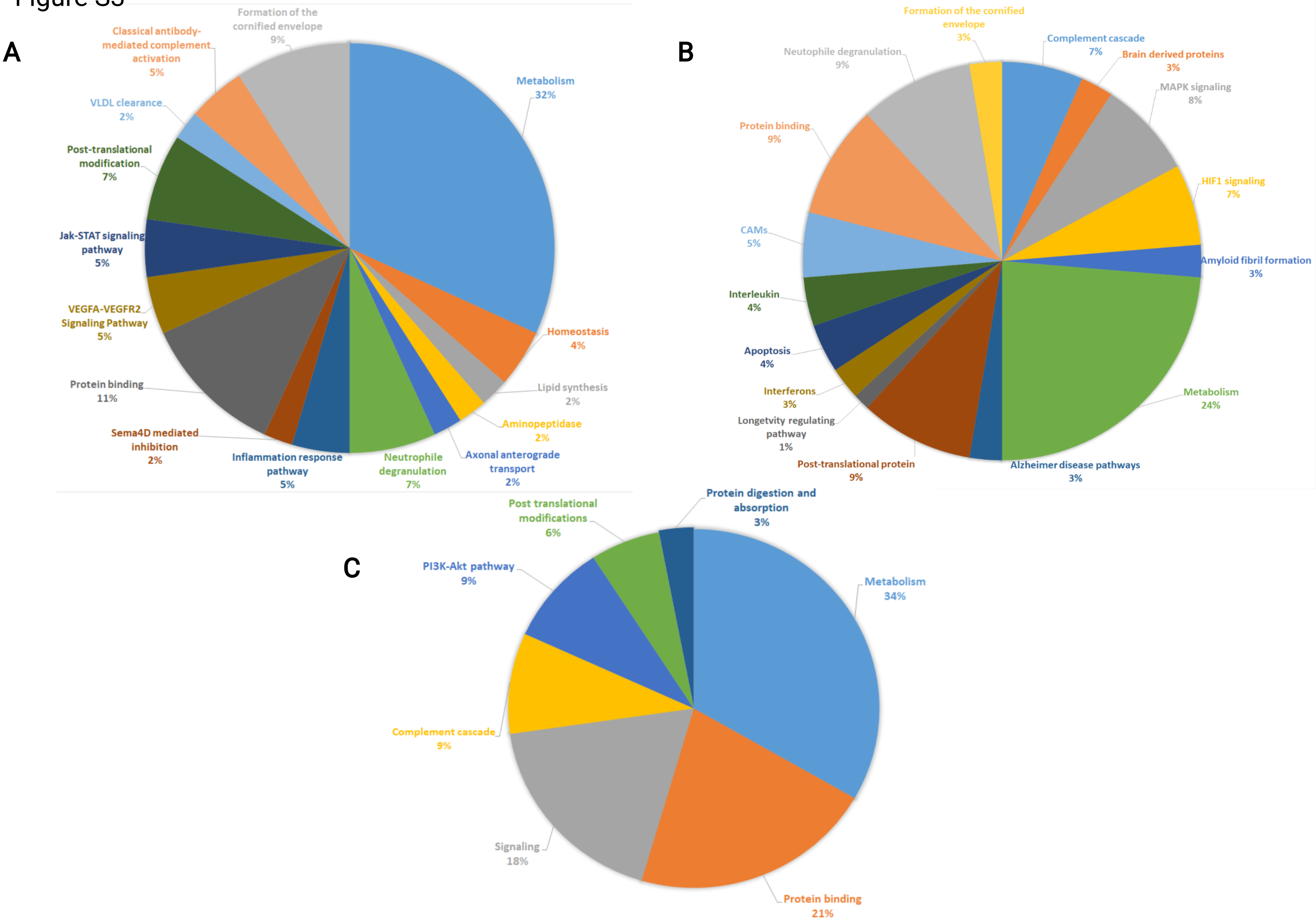

### Venn diagrams showed an overlap of 1,467 proteins in the three longitudinal comparisons

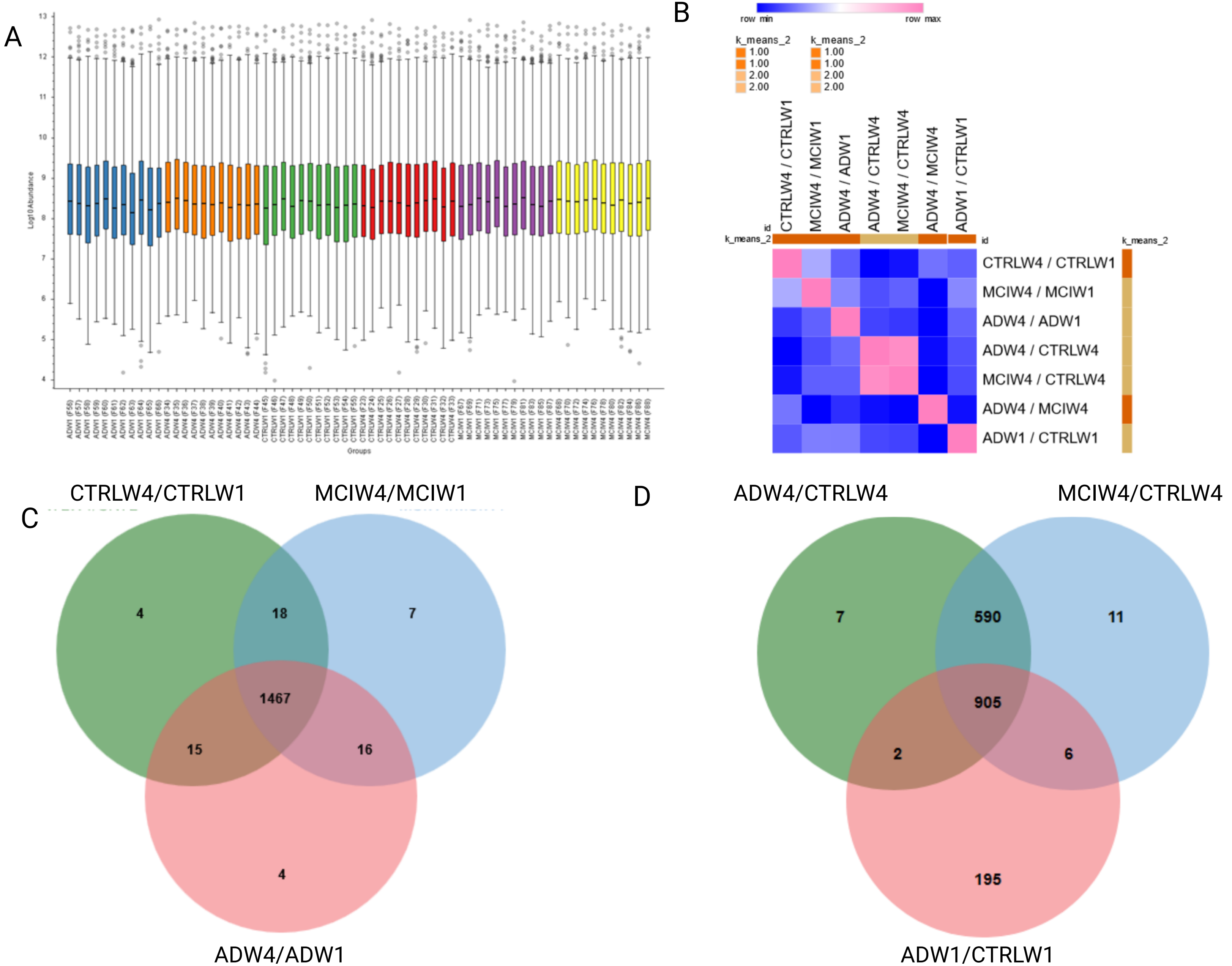
