## Supplementary material for "Deep proteome analysis of plasma reveals novel biomarkers of mild cognitive impairment and Alzheimer’s disease: A longitudinal study": Relatively few of the same DEPs were identified in both the cross-sectional and longitudinal analyses of AD and MCI, being 15 (Figure S6A and S6B) and

Figure S6

A

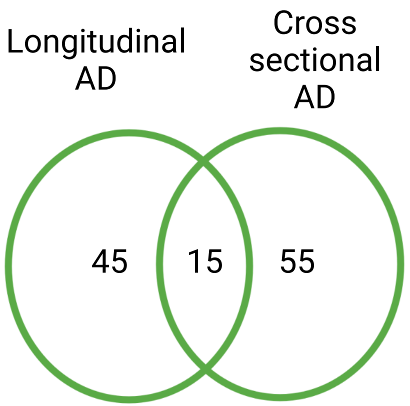


C

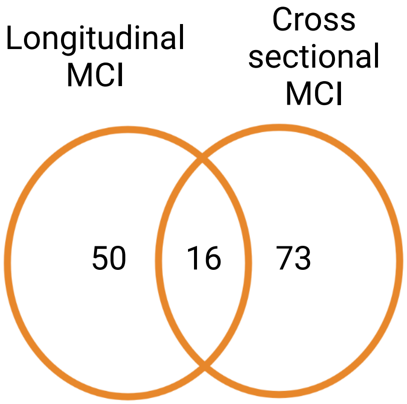


B

| Gene Symbol | PD2.4_Abundance Ratio: (ADW4) / (ADW1) | Scaffold_Fold change (ADW4) / (ADW1) | PD2.4_Abundance Ratio: (ADW4) / (CTRLW4) | Scaffold fold change (ADW4) / (CTRLW4) |
|---|---|---|---|---|
| EGFR | 0.679 | 0.2 | 0.525 | 0.6 |
| TF | **0.517** | 0.2 | 1.478 | 2.6 |
| ALDOB | 1.259 | 1.7 | 1.221 | 1.3 |
| QDPR | 1.334 | 5.4 | 1.342 | 1.8 |
| PAM | 1.372 | 1.5 | 1.27 | 1.4 |
| TNXB | 0.676 | 0.8 | 0.767 | 0.8 |
| LAMA2 | 1.249 | 1.4 | 1.202 | 2 |
| S100A7 | 2.049 | 3.5 | 1.749 | 1.5 |
| MAN2A2 | **0.56** | 0.2 | 0.706 | 0.4 |
| PSMB2 | 1.481 | 5.6 | 1.503 | 8.7 |
| VCP | 1.347 | 1.8 | 0.704 | 0.5 |
| CYCS | 1.369 | 6.6 | 0.679 | 0.7 |
| PTPRK | 2.076 | 2.3 | 1.322 | 2.2 |
| MAPRE2 | 1.639 | 3.7 | 0.598 | **0.3** |
| FAM3C | 1.555 | 2.8 | 1.409 | 2.4 |

D

| Gene Symbol | PD2.4_Abundance Ratio: (MCIW4) / (MCIW1) | Scaffold_Fold change (MCIW4) / (MCIW1) | PD2.4_Abundance Ratio: (MCIW4) / (CTRLW4) | Scaffold fold change: (MCIW4) / (CTRLW4) |
|---|---|---|---|---|
| PPBP | 2.159 | 1.2 | 0.84 | 0.6 |
| LTF | 1.288 | 1.6 | 1.242 | **1.5** |
| ASGR2 | 1.512 | 4.1 | 1.975 | 5.2 |
| PEPD | 0.758 | 0.8 | 0.685 | 0.8 |
| CPA1 | 0.71 | 0.2 | 0.692 | 0.2 |
| COL5A1 | 1.373 | 3.6 | 1.253 | 2.3 |
| IGFBP6 | 2.058 | **1.8** | 1.548 | 1.2 |
| HSPA4 | 2.136 | 3.9 | 2.037 | 2.7 |
| PSME1 | 0.748 | 0.5 | 0.717 | 0.2 |
| PPP1R7 | 1.777 | 2.9 | 1.678 | 2.1 |
| ADAMTSL4 | 1.415 | 1.3 | 1.268 | 1.3 |
| ITLN1 | 2.158 | 1.5 | 2.832 | 3.9 |
| C19orf10 | 2.555 | 4.3 | 1.293 | **5.2** |
| RARRES2 | 1.569 | 1.5 | 1.411 | 1.3 |
| FUCA2 | 0.689 | 0.5 | 0.804 | **0.3** |
| CECR1 | 0.723 | 0.6 | 0.654 | 0.5 |
